# Multiple dimensions of social motivation in adult female degus

**DOI:** 10.1101/2020.07.07.191486

**Authors:** Navdeep K. Lidhar, Ayushi Thakur, Anna-Julia David, Kaori Takehara-Nishiuchi, Nathan Insel

## Abstract

Many animals become more motivated to interact after a period of isolation. This phenomenon may involve general drives, e.g. for social touch or companionship, as well as drives that are specific to particular peers, and which ultimately serve to reestablish relationships between the individuals. Female degus are known to be affiliative with multiple other individuals, including unrelated and unfamiliar conspecifics, offering an opportunity to study social motivation independent from exclusive pair-bonds or overt, same-sex competition. We attempted to disentangle factors driving peer interaction by examining reunion behavior across several social isolation and separation manipulations. High levels of interaction were observed between adult females who had been separated even without isolation, revealing a drive to re-establish relationships with specific peers. The content of separation-only reunions differed from isolation, with the latter involving more early-session interaction, higher levels of allogrooming before rear-sniffing, and a higher ratio of chitter vocalizations. To assess whether post-isolation behavior was related to stress, we examined reunions following a non-social (footshock) stressor. Like isolation, footshock increased early-session interactions, but did not increase allogrooming before rear-sniffing or chittering, as compared with controls. To test whether separation-only reunion behavior shared qualities with relationship formation, we also examined reunions of new (stranger) dyads. Strangers exhibited higher levels of interaction than cagemates, with particularly high levels of late-session rear-sniffing. Like separation-only reunions, strangers showed more non-chitter vocalizations and lower levels of allogrooming before rear-sniffing. Across experiments, an exploratory clustering method was used to identify vocalizations that differed between conditions. This yielded promising leads for future investigation, including a chaff-type syllable that may have been more common during relationship renewal. Overall, results are consistent with the hypothesis that female degu reunions are supported by both general and peer-stimulus specific drives, expressed through the structure of physical and vocal interactions over time.

## Introduction

In many animal species, a period of social deprivation can cause a subsequent rebound of interaction. (e.g., humans [1] and rats [2]). The drive to interact after isolation is reminiscent of traditional models of motivation, in which the energy for a behavior increases as more time passes since the behavior’s expression [3, 4]. But social interaction can take different forms and satisfy different needs, many of which are conflated during isolation. One distinction that can be made is between “stimulus-specific” drives between individuals that have not seen one-another for a period of time and more general drives for, for example, social touch, companionship, or a sense of social agency. These two domains—stimulus-specific and stimulus-general—are difficult to isolate experimentally. In pair-bonding species, for example, general social drives are largely exclusive to a single individual, so the expression of motivation for one, specific peer is conflated with social motivation more generally. Some insight on stimulus-specific social drives comes from research on greeting behavior between individuals that have been separate for extended periods of time. Such greetings are thought to help reestablish dyadic relationships and trust [5–8]. Put another way, greetings reestablish predictive associations (expectations) that individuals have about one-other, including others’ probable actions and reactions, thereby reducing uncertainty and potentially promoting prosocial emotions and behavior. Complete social isolation, meanwhile, deprives an animal of all forms of social activity. The motivational impact of isolation may be at least partly mediated by neuroendocrine stress responses, as isolation causes an increase in systemic glucocorticoids [9] and social interaction, in turn, is known reduce or “buffer” stress levels [10–15]. A number of physiological changes are known to take place during social isolation [16–23], some of which can be directly linked with stress, and others that may be more independent. An important question is what contribution stress, by itself, contributes to interactive behavior after isolation.

While multiple different drives likely contribute to peer interaction after isolation, it remains unclear how each differentially impacts the content or structure of social interactions. The primary goal of the present study is to help build a “lexicon” of social behaviors associated with specific motivational contexts that can later be used to infer motivational states in less controlled settings. We focus our examination on the degu (*Octodon degus*), a caviomorph rodent from Chile. Degus are an interesting species for studies of social interaction on account of their high sociality [24], dynamic group structures [25–28], as well as the parallel research that can be done in both field and laboratory settings. Most importantly for the present experiments, adult female degus are highly interactive but do not show systematic preferences for familiar over unfamiliar peers [29]. This allows investigation of same-sex peer motivations without the confound of selective long-term bonds, meanwhile also minimizing aggressive behaviors that are typical between unfamiliar, same-sex adults of many rodent species. In native habitats, female degus are recognized for their constructive interactions, such as communal nursing, with both related and unrelated, same-sex individuals [26,30–33]. Degus also express a wide range of vocalizations in the human hearing range, and these have been manually classified based on spectral features and the social contexts in which they are used [34]. Degus therefore express a high dimensionality of social behavior that can be quantified to compare interactions between different experimental conditions.

We test four main hypotheses. Each experimental prediction assumes that more motivation results in more interactions during a 20 minute “reunion” session, and that different motivational drives are expressed with different interaction patterns in ways that generalize across dyads. Hypothesis 1: given that adult degus show high sociality in their natural habitat and in prior laboratory research, motivation to interact increases the longer they go without socializing. This predicts that animals will spend more time interacting following longer (24 hr) periods of isolation compared with very little (1 min) isolation time, with intermediate levels of interaction following short-term (45 min) isolation. Hypothesis 2: Degus form peer-specific relationships and/or predictive associations that require renewal after a period of separation. This predicts that A) a degu will become more motivated to interact with a peer following an extended (24 hr) period of separation, as compared with a “baseline” 1 min of isolation, and that B) separation-without-isolation reunions will differ from those following isolation. Hypothesis 3: Stress at least partly mediates increased interaction levels following isolation. This predicts that A) a non-social (footshock) stressor will also increase female social interaction and, B) reunions following footshock will share behavioral patterns with those observed following isolation. Finally, Hypothesis 4: adult female degus are highly motivated to interact with strangers due to drives that help establish new predictive associations and relationships. This predicts that A) stranger reunions will involve levels of interaction at least as high as those between reunited cagemates and B) there will be similarities between interaction patterns observed between strangers and those between separated-but-not-isolated cagemate dyads.

We record degu behavior during 20 minute reunions following manipulations that include social isolation, separating dyads without isolating them from other conspecifics, and exposure to an external stressor. The results reveal several subtle but notable distinctions in how different forms of motivation impact behavior.

## Methods

### Subjects

Thirty-eight degus aged 6 to 19 months (mean = 10.2 ± 3.6 SD) were used for all experiments. All degus came from our breeding colony at University of Toronto, descended from animals from University of Michigan and an Ontario breeder. By the start of the study the colony had been established in Toronto for approximately four years, and all degus used had been born in-house. Degus reach developmental maturity by 3.5 months [35, 36] and in the lab live up to 7 to 8 years [37]. We therefore consider the 6 to 19 month range as “young adult”. While the study was not designed to evaluate differences within this age range, a comparison of age against total interactive time yielded a negative result: n = 17, r = -0.22, p = 0.39) so age was not otherwise considered within the scope of the experiment. Prior to the start of experiments, degus were housed either in pairs within 14” x 11” x 9.5” or, in the case of Experiment 2, in groups of four within 15” x 19” x 8” polycarbonate cages. All were housed in a vivarium at the biological science facility at the University of Toronto. Paired and group housing was usually with siblings, but in Experiment 1, three of the nine pairs were non-sibling (placed together sometime between weening and one month prior to the experiment). Prior research has found that, in degus, familiarity more strongly determines social interactions than kinship [38]; but effect sizes are described and compared for sibling and non-sibling pairs in Results. the Degus were kept on a 12:12h light/dark cycle with food and water available ad libitum. All procedures were conducted in accordance with the policies of the biological facilities at the University of Toronto and all protocols were approved by the University of Toronto Animal Care Committee.

### Apparatuses

Behavioral recording took place in degus’ homecages placed inside a 15’’x19’’x 24’’plexiglass chamber. The ceiling of the recording chamber was embedded with a Sony (Sony; Tokyo, Japan) USB2 camera (video sampled at 30 Hz) and Sony lapel microphone (model number ECMCS3, sampled at 44100 Hz). Data were collected using Bandicam multi-media recording software (Bandisoft; Seoul, Korea). Social isolation was peformed by placing each degu into separate, 14” x 11” x 9.5” chambers, either in the experiment room (for 1 and 45 min isolation periods) or the vivarium (for 24 hr isolations). Footshocks used for the “acute stress” condition in Experiment 3 were delivered in a 18” × 17.5” × 15” conditioning chamber with a metal grid floor consisting of 5 mm thick stainless steel rods spaced 1 cm apart (Colbourne instruments; Holliston, USA).

### Protocol

Habituation: Prior to behavioral testing, degus were acclimated to the recording chamber. Degu pairs were brought to the conditioning room and their homecage was placed into the soundproof recording chamber. The pair spent 1 hour undisturbed in the chamber for each of 3 consecutive days. Experimental protocols began immediately after acclimation except in the case of Experiment 4, for which acclimation and prior experimentation took place at varying times prior to the reunion.

Experiment 1: To test the effects of social isolation on interactive behavior, 9 pairs of female cagemate degus (18 degus) were placed in separate cages for either 1 or 45 min in the recording room, or 24 hr in the vivarium (the difference in holding room was logistical, as the shorter intervals provided insufficient time for experimenters to shuttle both degus back to the vivarium, and animals were required to be in the vivarium overnight). The isolation chambers were the same size as the animals’ homecages and were transparent and contained clean bedding. Notably, although they were “isolated” in the sense of being unable to touch or interact with other individuals, they could still see and hear other degus through the cage. Members of a dyad were kept on different sides of the room to prevent direct communication between them. Following the social isolation period, the original homecage was placed within the soundproof chamber for 20 minutes, during which time video and audio recordings were collected. The basic protocol is illustrated in **Fig 1A**.

**Fig 1.**
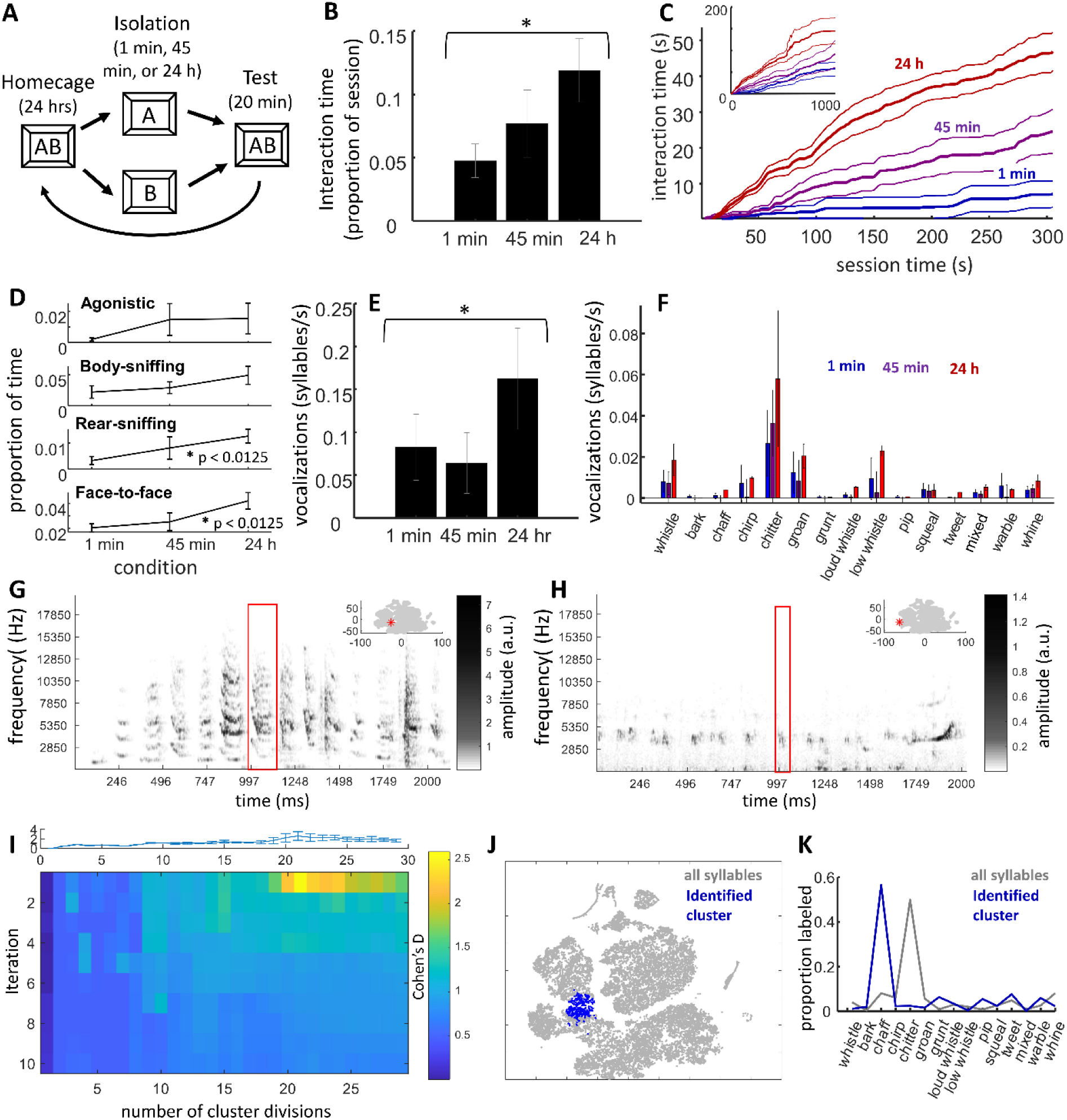
Effects of social isolation time on reunion behavior. **A)** Illustration of experimental design. Degus (n = 9 dyads) were separated from cagemates for 1 min, 45 min, or 24 hours then interactions were examined in a testing chamber. After being reunited for 24 hours the process was repeated. **B)** Proportion of time engaged in physical interactions (y axis) increased with the amount of preceding time of isolation (x-axis). **C)** Cumulative distributions of physical interactions over session time. Thicker lines are means, thin lines s.e.m. Higher levels of interaction in the 24 hr condition were seen within the first minute and remained high for the first 5 minutes. Inset shows full, 20 min of session time; although interaction time in the 24 hr condition remained higher, interactions in the 1 and 45 min conditions were more distributed across session time. **D)** Line graphs of interaction time across types of physical interaction; no interaction between type and condition was observed. **E)** Vocalization rates (y-axis) significantly increased across conditions (x-axis). **F)** Vocalization rates were consistently higher in the 24 hr condition across the 15 vocalization categories tested. **G)** Example spectrogram of a series of chaff and related syllables and **H)** chitter syllables. Low-frequency (below 400 Hz) noise is removed from plot. Red box in the center highlights a single syllable, inset in upper-right shows the t-SNE map used in panel J and **S3**, with a red asterisk marking where the central syllable landed within the projection. Syllables in both sequences may have come from one or both members of a dyad. **I)** Heatmap of identified syllable clusters with the strongest difference between the 24 hr and 1 min conditions, across k-means clusters repeated for 2-30 cluster divisions (columns), iterated after removing syllables from the best cluster (rows). Plot on top is an errorbar showing variance of the first (top-row) across 20 repetitions. The hot-spot in the heat map shows that when syllables were divided into 20 clusters, one cluster, consistent across the 20 repetitions, showed a particularly high effect size and was significantly, across dyads, more highly expressed in the 24 hr condition. **J)** Map of t-SNE projection of vocalization features. The identified cluster on this projection is represented in blue dots. **K)** Proportion of manual label types across syllables across all studies (grey) is plotted alongside the proportion of labels for the identified cluster. Syllables in this cluster were predominately labeled as “chaffs”, and almost never as “chitters”.

Experiment 2: To test the differences between social isolation and separation-without-isolation, each of 4 groups of 4 cagemate degus (16 degus) were first separated into two cages of only two degus (SEP). Following 24 hours, one member of each of the new cages was placed together in a clean 14” x 11” x 9.5” cage within in the recording chamber for a 20 min reunion session, after which the recorded pair was taken to the vivarium and kept together in that same cage for the subsequent 24 hours. The same was done with the remaining members of the two cages. Following 24 hours as reunited cagemates, the dyad was separated into two individual cages for a 1 minute period and then returned for a second, 20 minute reunion (CTL). Degus were then returned to separate cages and remained isolated for 24 hours, after which a final, 20 min reunion session was performed (ISO).

Experiment 3: To test the impact of stress on social interactions, the same degus from Experiment 1 were presented with two additional conditions. In one, cagemate dyads were separated into two different cages in two separate experiment rooms for a 45 minute period. During this time, following a 5 minute adaptation period, 8 sunflower seeds (a familiar, preferred food) were placed one at a time over a 2.5 minute period in each cage (RWD). When the 45 minute period had elapsed, they were placed together in their homecage, in the recording chamber for a 20 minute reunion session. In the other condition, cagemate dyads were separated into two separate rooms and placed in a conditioning chamber with a metal grid. Following a 5 minute baseline period, degus received 10, 1.0 mA footshocks lasting 2 s each over a 2.5 minute period (SHK). This shock level and pattern was based on a protocol we used previously to evoke contextual and observational fear learning [39], though to reduce total aversiveness we decreased the number of shocks by one-half. The 1 mA level offered a compromise for maximizing animal welfare while also eliciting a sufficient stress response (in rats, a series of 1.5 mA footshock stimulations evokes significantly higher levels and longer-lasting glucocorticoid responses compared with 0.5 mA [40]). After receiving footshock, degus were placed in separate cages for the remainder of the 45 minutes.

To ensure that the reward and stress conditions did not contaminate subsequent exposures, degus were always tested with the neutral (NTL) 45 minute condition first (part of the Experiment 1), then RWD, and then SHK. Ultimately, no evidence of progressive behavioral changes across these conditions were observed, though issues of order effects are taken-up more thoroughly in Discussion (*Overview & limitations*).

Experiment 4: To investigate how interactive behavior was impacted by social novelty, potentially associated with relationship establishment, degus were united with unfamiliar, stranger conspecifics following 24 hours of isolation. Animals used for Experiment 4 were drawn from those from Experiments 2 (12 degus) or were members of a separate set of animals that had received pre-exposures and were subsequently used for a separate study (4 degus). Degus that had been involved in another experiment had been pair-housed with a cagemate for at least one week after the prior experiment had ended.

The sequence of reunion sessions for each dyad are illustrated in Table 1. Notably, the order of conditions was not fully counterbalanced, and the potential implications of that confound are taken-up in Discussion (*Overview & limitations*).

**Table 1.** Sequence of reunion exposures for each dyad. Conditions are labeled as Experiment number – condition (SEP = separation, CTL = 1 min control, ISO = 24 hr isolation, RWD = reward, SHK = footshock, STR = stranger). Numbers reflect the order that the dyad experienced the condition. Blacked-out cells show reunion conditions that a dyad did not participate in. The 45 min condition used in Experiment 1 was re-used as the neutral control in experiment 3, involving the same dyads.

| Dyad | 1-1min | 1-45min / 3-NTL | 1-24hr | 2-SEP | 2-CTL | 2-ISO | 3-RWD | 3-SHK | 4-STR |
| --- | --- | --- | --- | --- | --- | --- | --- | --- | --- |
| 1 | 3 | 2 | 1 |  |  |  | 4 | 5 |  |
| 2 | 1 | 3 | 2 |  |  |  | 4 | 5 |  |
| 3 | 1 | 3 | 2 |  |  |  | 4 | 5 |  |
| 4 | 1 | 2 | 3 |  |  |  | 4 | 5 |  |
| 5 | 1 | 2 | 3 |  |  |  | 4 | 5 |  |
| 6 | 1 | 2 | 3 |  |  |  | 4 | 5 |  |
| 7 | 1 | 2 | 3 |  |  |  | 4 | 5 |  |
| 8 | 1 | 2 | 3 |  |  |  | 4 | 5 |  |
| 9 | 1 | 2 | 3 |  |  |  | 4 | 5 |  |
| 10 |  |  |  | 1 | 2 | 3 |  |  |  |
| 11 |  |  |  | 1 | 2 | 3 |  |  |  |
| 12 |  |  |  | 1 | 2 | 3 |  |  | 4 |
| 13 |  |  |  | 1 | 2 | 3 |  |  | 4 |
| 14 |  |  |  | 1 | 2 | 3 |  |  | 4 |
| 15 |  |  |  | 1 | 2 | 3 |  |  | 4 |
| 16 |  |  |  | 1 | 2 | 3 |  |  | 4 |
| 17 |  |  |  | 1 | 2 | 3 |  |  | 4 |
| 18 |  |  |  |  |  |  |  |  | 1 |
| 19 |  |  |  |  |  |  |  |  | 1 |

### Behavioral analysis

Software: Analysis relied on several software packages. Scoring of physical behavior was performed using BORIS (Behavioral Observation Interactive Research Software; Torino, Italy [41]), which allows users to log events during video playback. Scoring of vocalization behavior was performed using Audacity (Audacity v.2.1.2; Pittsburg, USA), as well as Sound Analysis Pro (SAP), a vocalization detecting software developed for segmenting syllables and vocalization structure in birdsong [42]. All other analyses were custom-written in MATLAB, as described below and provided at https://github.com/ninsel/DeguReunionMotivation.

Physical interactions: Each video recording was manually scored by blind raters using BORIS. Blinding was done by using coded filenames. Ethograms included four general types: *agonistic encounters* (bites, characterized by face-to-body contact of one degu with the other making a startled movement away; mounting, involving one degu placing forepaws on the other while the other made avoidance movements; wrestling, characterized by rapid movements of both bodies and usually accompanied by other aggressive actions; punching/boxing, with degus sometimes rearing on hindlegs; body-slams, in which degus kicked with their hind legs, and any other apparently aggressive encounter), *allogrooming/body sniffing* (one degu’s nose touching and/or interacting with the neck or body of the other), *rear-sniffing* (a.k.a. anogenital investigation, where one degu’s nose was near and directed toward the base of the tail of the other), and *face-to-face* interactions (one degu’s nose near and directed toward the nose or mouth of the other). Ethograms were created based on a relatively minimal set of highly distinguishable behaviors that were observed across most sessions, with a goal of maximizing simplicity and inter-rater reliability. To assess inter-rater reliability, several sessions were scored by more than one rater, and Cronbach’s alpha was calculated to be 0.97 (very high). For each scored behavior, the identity of the animal was also included. Often behaviors overlapped: e.g., one degu may initiate rear-sniffing and the other may then reciprocate before the first behavior ended. In the case of face-to-face, the action was necessarily mutual, but these were still labeled with the identity of the degu that initiated the interaction. One potential problem with behaviors overlapping in time is that the sum of all behaviors performed by one degu added to the sum of all behaviors performed by the other could, in principle, exceed the total time of the session. Analyses therefore combined interaction time from each degu by considering all time during which *either* one degu *or* the other was engaging in the action (i.e., the union of time, rather than the combined total of both).

Comparisons of physical interactions between experimental conditions were analyzed using the following three tests: 1) Total time dyads spent interacting, and total time engaging in different interaction types. As noted above, interaction time was primarily analyzed as all time that either one or the other degu of a dyad was engaging in an interactive behavior (body-sniffing, face-to-face, etc.). 2) Temporal distribution of interactions across the session, primarily quantified and illustrated by cumulative distributions of interaction over time. Cumulative distributions of interactions were computed with a 2 s bin for each session (discussed below in *Statistics*). 3) Temporal relationships between interaction types, quantified and illustrated by the cross correlograms. Cross correlograms evaluated the time-lagged coincidence of one interaction relative to another. To compute these, a histogram was generated: times preceding zero (left side of the x-axis) showed the incidence of one (“target”) action relative to when another (“reference”) action was initiated, while times following zero (right side of the x-axis) showed the incidence of the target action relative to when the reference action had terminated. Actions that took place simultaneously were therefore not included in the histogram. These histograms were then normalized by the number of reference actions, thereby converting incidence into rates. For interpretability, the histograms were further normalized by dividing by the proportion of time during the session that the target action took place. In other words, values of “1” in the cross correlation meant there was no observable temporal relationship between the two interaction types. Importantly, this means that y-axis units were not correlations coefficients, but normalized rates. To assess significance, all values from -10 to -1 s (before onset) and from 1 s to 10 s (after offset) were averaged within each session, and a paired t-test was performed to assess whether values were A) consistently higher than 1, and whether values were B) consistently higher than the average of ten surrogate datasets, in which onset/offset times of the comparison action type were randomly shuffled within a -5 to 5 minute interval. Only those cross correlations that met both criteria at an alpha value of 0.05 were considered significant. In cases where comparisons were being made between conditions, a paired or two-sample t-test was used. We address the issue of multiple comparisons below under the *Statistics* subheading.

Vocal interactions: Vocalizations were scored by loading audio files into Audacity and using both audio and visual spectrograms to identify vocalization start and end times. Blinding of vocalization scoring used the same coded filenames as used for coding physical interactions. To reduce noise, the Audacity “noise reduction” feature was used, which creates a filter based on a selected audio baseline period (baseline was several seconds of audio recorded prior to degus placed in the recording chamber). Experimenters scored vocalization by loading individual sound clips generated by Audacity into SAP, where spectrograms and summary statistics (amplitude, frequency modulation, duration) were compared with those presented by Long (2008; [34]). Inter-rater reliability was estimated using Cronbach’s alpha = 0.96 (very high). Comparisons of vocal behavior between sessions relied on the following tests: 1) total vocalization rates (number of syllables over session time), and rates of specific, manually scored vocalization types, and 2) differences in the profile of vocalization types detected through unsupervised clustering of the syllables (repeated-k-means).

We chose to use an unsupervised clustering method over a supervised algorithm out of interest for a method that was complimentary to, rather than dependent on, manual-labeling. Unsupervised methods have the potential of identifying syllable clusters based on features that human listeners are not as well tuned to, thereby revealing syllable types that we would not have known to look for. The sequence of steps involved in this analysis consisted of the following: 1) First, a set of 31 features was computed for each identified syllable. The first feature was syllable duration, while the remaining thirty were dominant frequency, spectral dispersion, and amplitude across each of ten deciles of time. These values were standardized across all syllables by converting each of the 31 features to a z-score relative to all other recorded vocalizations, and then weighted (each multiplied by a scalar) such that syllable duration counted for 10 times each dominant frequency value, 20 times each spectral dispersion value, and 40 times each amplitude value. The weightings were chosen based on subjective assessment of which features should play the strongest roles in syllable classification, an approach that has precedence in the field [43]. 2) To reduce the dimensionality of the feature space, and also account for correlations between features (e.g., adjacent deciles are likely to be correlated), we converted features to principle components. 3) Describing each syllable by its first 10 principle components (accounting for 99.5% of the variance), we then entered all recorded syllables across all experiments into a k-means clustering method. K-means clustering was used to subdivide the pool into 2 clusters, 3 clusters, 4, and on up to 30 clusters (10 replicates each, with 20 repetitions each to assess cluster stability). 4) For each syllable cluster, the number of cases in each session was identified. The proportional rate of syllables in one condition could then be compared with another condition by computing an effect size (Cohen’s D) and a significance value (p value from a paired t-test when considering the same animals, or two-sample t-test in Experiment 4 when comparing different dyads). In some cases, more than one cluster might show a strong difference, with some being sub-divisions of others. To identify only the *independent* groups of syllables that differed between conditions, only the strongest cluster was considered, then all syllables from that cluster were removed and significance testing between clusters was performed again, across multiple iterations until no significant clusters were identified. This k-means clustering method was modified from a similar method used in previous work from our group, which was designed to evaluate whether there were subdivisions of distress/alarm calls [39]. The previous version differed in only minor respects; e.g., only four features were included, so no dimension reduction was needed, and notable clusters were identified with their p-values rather than effect sizes. Additionally, since there was expected to be fewer categories, the earlier version did not repeated the process iteratively after removing syllables of the strongest category.

One acknowledged risk of the repeated-k-means analysis is that if syllables are randomly distributed across all sessions, one might expect to observe significant clusters roughly 5% of the time, given an alpha of 0.05. Empirically this was not the case; however, taking this concern into account, identification of clusters was treated as purely exploratory. For partial validation of the results, we performed an empirical assessment of the individual clusters (including consistency of the cluster across the 20 repetitions, as measured by the Rand index, and how the cluster mapped onto manually-labeled vocalizations) and present figures of effect size distributions for qualitative comparisons between experiments. All identified clusters were reported in Results, and to reduce biases from multiple-comparisons across repetitions only the median significance values and effect sizes across the 20 repetitions were reported.

Although behaviors were performed within a soundproof chamber, it was still possible that vocalizations below a certain volume were undetected by out methods. To assess whether this might be the case, vocalization syllables were segmented into 10 ms bins and the peak amplitude was measured relative to the dominant frequency at that amplitude. This was first done across syllables recorded in several, adult female degu “reunion” experiments which used the same methods (24,535 syllables). An initial look at the data revealed that a substantial proportion of syllables (approximately 43%) were not loud enough, or of high enough frequency, to exceed the volume of the ambient noise at 400 Hz (the lower cutoff that was used to compute dominant frequencies). Syllables were therefore separated into two populations: those with amplitude peaks that exceeded a 400 Hz cutoff, and those that did not. Amplitudes and dominant frequencies of those that reached threshold are illustrated in **S1A**; of these, only one small grouping of low-amplitude, low-frequency syllables, almost exclusively labeled as chitters, appeared to have a distribution with the lower-end cut-off from detection. Of those that did not stand out from the noise, also most often labeled as chitters, the distribution also appeared to be positively skewed (**S1B**). These data suggest that only a minority of vocalizations failed to reach detection threshold, and most of these were likely to be “chitters.”

### Statistics

Parametric statistics are used in nearly all comparisons, including repeated measures ANOVAs (used for all one-way tests), two-way ANOVAs (to evaluate interaction effects), and both paired (for same-dyad comparisons) and two-sample (for between-dyad comparisons) t-tests. In the case of manually scored vocalizations, variance across dyads was not distributed normally and therefore a Wilcoxon rank sum test was used. Generally, sample sizes were low, and the high probability of false negatives is addressed in discussion.

Many analyses addressed specific predictions in a given experiment; however, there were several circumstances in which multiple comparisons were used in an exploratory fashion, raising the possibility that false positives may arise by chance. The first of these is when testing whether each individual behavior (agonistic, grooming/body-sniffing, rear-sniffing, face-to-face) is affected by conditions within an experiment. In this case, we employ a Bonferroni correction when assessing statistical significance (alpha = 0.05/4 = 0.0125). The second circumstance is when looking at temporal interactions for each relationship between behaviors (e.g., rear-sniff before grooming, rear-sniff after grooming, etc). Here, cross correlations had to reach two criteria (changes from baseline and differences from a surrogate dataset) to be considered significant. The analysis is performed with acknowledgment of a false positive rate of up to 5%, though most comparisons elicited much lower p-values, and causal inferences focus on those relationships that showed consistent and meaningful patterns between experiments. Finally, multiple comparisons are used when assessing differences between conditions over different, cumulative time intervals of the reunion session. In this case, a t-tests was used to compare each of 600, 2 s intervals between conditions. This method was selected as an exploratory approach that made no *a priori* assumptions on the distribution of behaviors over time, or how that may vary between conditions (as a contrast to model-fitting, which would require would require either overfitting a high number of parameters or assumptions about the distributions). To best address the possibility of false positives, only extended intervals are reported, along with session times when differences reach their peak, and the method is complement with a secondary, directed analysis comparing the slopes of the cumulative distributions.

## Results

### Experiment I: Isolation time

We first sought to confirm that degus become increasingly motivated to interact following increasing intervals of social isolation. Nine, adult female degu cagemate dyads were pre-exposed to a behavioral recording chamber for one hour each day over three days and then, on sequential days, isolated from one another in separate cages for 1 min, 45 min, and 24 hr (**Fig 1A**). Following each isolation period, dyads were brought back together for 20 minutes in the recording chamber while video and audio data were collected. Consistent with expectations, there was a significant impact of isolation time on the time degus spent physically interacting (repeated-measures ANOVA, F_2,16_ = 4.93, p = 0.021, **Fig 1B**). The effect of isolation was particularly strong earlier in the session, generally peaking around 3 to 4 min (**Fig 1C**; using an alpha of 0.05, differences between 1 and 45 min conditions were observed for the entire interval between 78 and 422 s, peaking at 224 s: t_8_ = 4.79, p = 0.0014; differences between 24 hr and 45 min observed for the entire interval between 30 s and 1198 s, peaking at 282 s, t_8_ = 9.48, p = 1.26 x 10^-5^; and differences between 24 hr and 1 min persisted throughout the session, significant by 20 s and peaking at 194 s, t_8_ = 10.30, p = 6.81 x 10^-6^). Interactions were parsed into four types: agonistic (which included mounting), allogrooming/body sniffing, rear-sniffing (i.e., anogenital investigation), and face-to-face. No interaction was observed between condition and interaction type (2-way ANOVA, F_6,96_ = 0.53, p = 0.78). Examining each interaction type individually, significant differences in interaction time (alpha = 0.05/4 = 0.0125, Bonferroni correction) were observed for rear-sniffing (F_2,16_ = 5.9, p = 0.012) and face-to-face interactions (F_2,16_ = 6.78, p = 0.0074), with a trend also observed for grooming/body sniffing (F_2,16_ = 3.58, p = 0.052; **Fig 1D**). The time spent in agonistic behavior did not appear to vary across the conditions (F_2,16_ = 0.23, p = 0.80).

When degus were not interacting with one-another, they were typically rearing and looking upward along the walls or sniffing and sometimes digging into the chamber bedding. Self-grooming was the only non-interactive behavior systematically scored, which significantly increase with preceding isolation time (F_2,16_ = 4.95, p = 0.021). Although the study was not designed to compare behavior in sibling versus non-sibling cagemates, there appeared to be no meaningful difference in the effect size of the 24 hr versus 1 min isolation in non-siblings (n = 3, Cohen’s D = 0.95) compared with siblings (n = 6, Cohen’s D = 0.82).

To assess the structure of behavior, we examined whether there were temporal relationships between physical behaviors. We had no a priori hypotheses on how patterns might vary with increasing isolation time, and anticipated that lower levels of interaction in short (1 min) interaction conditions would impact the validity of a direct comparison between conditions. Initial analyses therefore pooled all interactions across the 1 min, 45 min, and 24 reunion conditions to identify a base-set of relationships, providing a framework for interpreting interactions observed in subsequent experiments. Cross-correlograms were generated for each pair of interaction types (e.g., face-to-face and rear-sniffing, rear-sniffing and allogrooming/body sniffing, etc), where values on the negative side of the x-axis (time) represented the occurrence of an action type *before the onset* of another “reference” type, and values on the positive side of the x-axis represented the occurrence of an action type *following the offset* of the reference type. A significant temporal relationship was determined if the cross correlogram values were A) consistently higher than baseline (i.e., systematically higher than 1), and B) consistently higher than surrogate datasets, in which onset/offset times of the comparison action were randomly shuffled within a -5 to 5 minute interval (see Methods). In the initial analysis performed on Experiment 1 data Significant relationships (described as comparison relative to reference) were identified for: 1) allogrooming/body sniffing preceding rear-sniffing (above baseline: t_16_ = 2.84, p = 0.010, above shuffled: t_16_ = 4.12, p = 5.0 x 10^-4^), 2) agonistic preceding and following agonistic (above baseline: t_16_ = 2.49, p = 0.024 and t_16_ = 3.18, p = 0.0059 respectively, above shuffled: t_16_ = 2.59, p = 0.020 and t_16_ = 3.33, p = 0.0042), 3) rear-sniffing following allogrooming/body sniffing (above baseline: t_16_ = 2.45, p = 0.0024, above shuffled: t_16_ = 5.68, p = 0.0010), 4) allogrooming/body sniffing following face-to-face (above baseline: t_16_ = 3.40, p = 0.0023, above shuffled: t_16_ = 4.13, p = 4.0 x 10^-4^) 5) face-to-face preceding and following allogrooming/body sniffing (above baseline: t_16_ = 4.65, p = 1.0 x 10^-4^, t_16_ = 2.52, p = 0.018 respectively, above shuffled: t_16_ = 5.88, p = 3.94 x 10-6, t_16_ = 4.16, p = 3.0 x 10^-4^). Averaged cross correlograms are presented in **S2**, are specific examples in subsequent sections.

As with physical interactions, isolation also appeared to increase vocalizing, measured by the frequency of syllables over session time (F_2,16_ = 4.18, p = 0.034, **Fig 1E**). To unpack this result, vocalization syllables were manually and automatically parsed into types. Manual classification included 32 categories, guided by previous research on degu social vocalizations[34]. Of these, only approximately 14 vocalization types were observed at least once, on average, per session— for the purposes of analysis, all others were included within a 15^th^ “mixed” category. Each vocalization type showed high variability between dyads and sessions, and no significant interaction was observed between condition and vocalization type (2-way ANOVA, effect of condition: F_2,360_ = 3.04, p = 0.049, effect of type: F_14,360_ = 5.22, p = 5.3 x 10^-9^; condition x type, F_28,360_ = 0.38, p = 1.00, **Fig 1F**). Examples of sequences of chaff- and chitter-scored vocalizations are illustrated in **Fig 1G** and **1H** respectively, with examples across categories presented in **S3**.

Differences between manually-scored syllable categories were sometimes subtle, and there was likely discrimination ambiguity between certain types (illustrated by the overlap of the different classes in feature space, **S4;** also confirmed by post-hoc inspection of syllables). We therefore also used an unsupervised clustering method to assess whether clusters of syllables with shared features might be differentially expressed between conditions. The repeated-k-means approach is detailed in Methods. Briefly: syllables were divided into clusters using a k-means algorithm (2 to 29 clusters, repeated 20 times, all across 10 iterations of removing the best cluster from the previous) and syllable clusters that made-up a higher proportion of those expressed during one condition compared with another were identified. For the purposes of this investigation, consistency across dyads, was emphasized over magnitude of difference, and proportion of syllable type within a session emphasized over total numbers. The method was considered an exploratory complement to manual labeling that, aside from syllable start and end times, was unbiased by the experimenter.

Comparing the 24 hr to 1 min isolation conditions, only one cluster stood out as showing a systematic (across dyad) difference (**Fig 1G & H**). This group of syllables was most precisely isolated when the pool was divided into 20 clusters (median p across repetitions = 0.013, median Cohen’s D = 2.7; consistency of cluster across iterations was also high: average adjusted random index = 0.96, where 1 is perfect cluster overlap). This syllable type was often manually scored as a “chaff” (**Fig 1I**) and was distinguished by relatively high average dominant frequencies around 6000 Hz, slightly lower-than-average entropy or frequency dispersion (i.e., more “scratchy”) and slightly higher than average amplitude. These vocalizations were not more temporally associated with specific interactive behaviors than average vocalizations, although prior work has suggested that chaffs may be more agonistic in nature [34].

To summarize the findings of Experiment 1, we first confirmed our prediction that adult, female degu dyads interact with one-another more following increased isolation time. The increased interaction did not significantly differ across interaction types, though notably there was no sign of increases in agonistic behavior. Pooling conditions together, we found a number of significant temporal interactions between interaction types, providing a starting point for more directed analyses of behavioral structure in future experiments and studies. Vocalizing also increased with longer isolation times. While differential increases across vocalizations were not observed with increased isolation time, this could have been impacted by low power, and exploratory analyses did identify one cluster of chaff-type syllables that might have been differentially increased in the 24 hr condition.

### Experiment II: Isolation vs. Separation

The second experiment examined whether dyads interact differently when motivated by prior social isolation, as compared with prior separation from one-another without being isolated (i.e., “general” versus “stimulus specific” social motivators). This was tested by comparing interactions in reunited dyads following three manipulations (illustrated in **Fig 2A**): 1) Cages of four female degus were split into two cages for 24 hours, thereby causing separation of two dyads (SEP); 2) all individuals were socially isolated for 24 hours (ISO); and 3) all individuals were isolated for only a 1 min interval (CTL).

**Fig 2.**
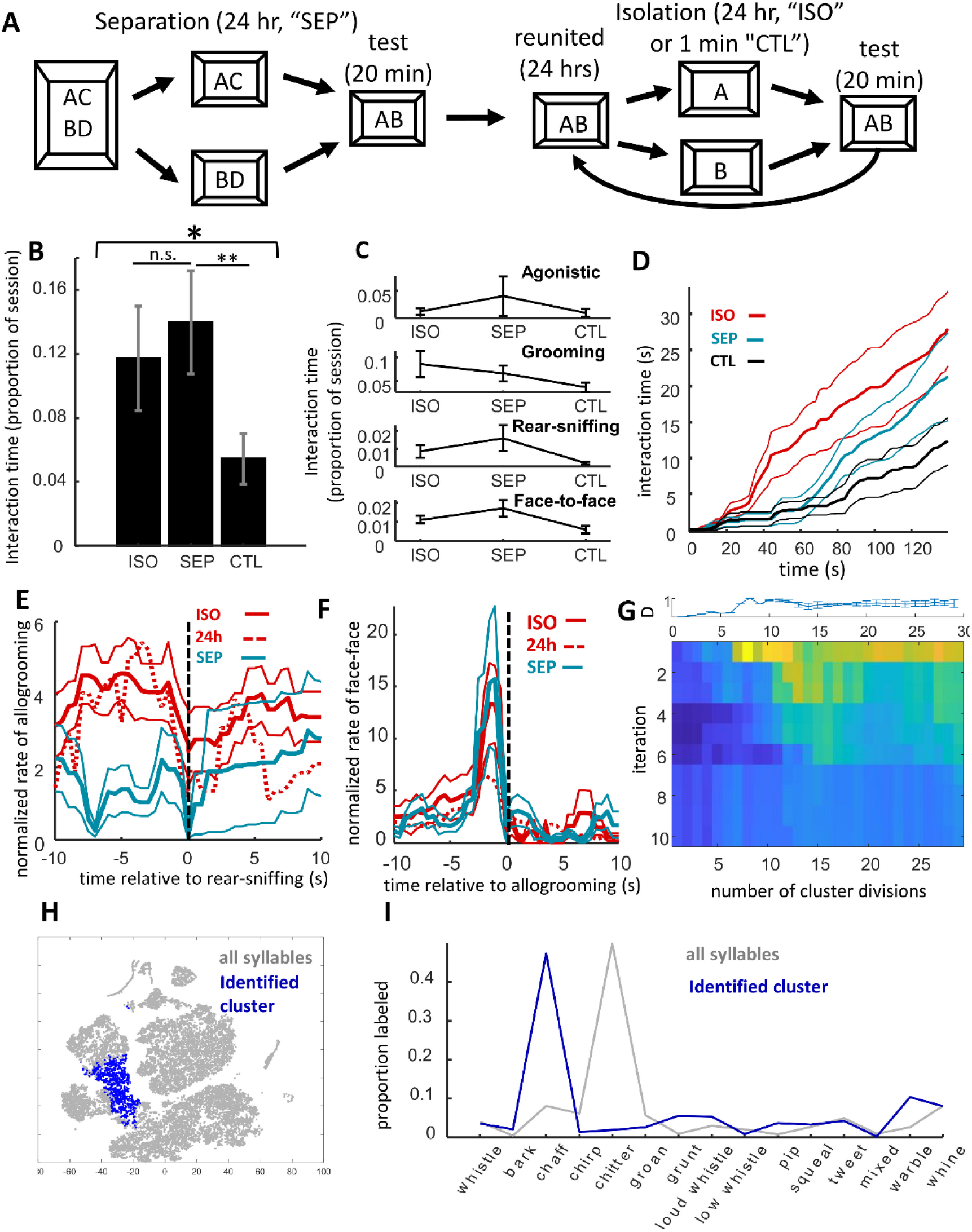
Comparison of isolation versus separation on reunion behavior. **A)** Illustration of experimental design. Four cages of four (n = 8 dyads) were split into two cages for 24 hours, separated dyads were then united for reunion SEP tests. Subsequently these dyads were isolated for 1 min and then 24 hours for CTL and ISO tests respectively, reunited for 24 hrs between each. **B)** Proportion of time degus engaged in physical interactions was at least as high in SEP as ISO, but lower in CTL. **C)** Interaction time for different types of physical interactions. There were no systematic differences between SEP and ISO, though patterns hint at higher rear-sniffing and lower grooming in SEP relative to ISO. No between-group comparison met the criteria for significance (alpha = 0.0125). **D)** Cumulative distribution of physical interactions over session time (thicker lines are means, thin lines s.e.m.). Although interactions in SEP were high over the 20 min, following isolation degus were much more likely to interact during the first minute of the reunion session. **E)** Cross correlogram of levels of allogrooming/body-sniffing (traces) relative to rear-sniffing. Negative time values are prior to rear-sniffing initiation, positive are after termination of the behavior. Both ISO and the 24h condition from Experiment 1 showed higher allogrooming before rear-sniffing compared with SEP. **F)** Cross correlogram of the levels of face-to-face interactions relative to allogrooming/body sniffing. A significant increase in face-to-face contact before allogrooming was observed for all conditions. **G)** Heatmap of identified syllable clusters with the strongest difference between ISO and SEP, with increasing cluster divisions across columns and iterations after removing syllables from the best cluster down rows. Plot on top is an error bar showing the variance the first (top-row) across 20 repetitions. **H)** Map of t-SNE projection of vocalization features. One cluster was identified of syllables that differentiated between SEP and ISO conditions (blue points). **I)** Proportion of manual labeled types across all syllables (gray trace) and for only those syllables in the identified cluster (blue trace) show that, as with the cluster identified in Experiment 1, these were chaff-type syllables.

Degus interacted more in SEP than they did in CTL (repeated-measures ANOVA effect of condition F_2,14_ = 4.93, p = 0.024; SEP vs. CTL, paired t-test, t_7_ = 3.94, p = 0.0056; **Fig 2B**), and there was no detectable difference between ISO and SEP (t_7_ = 0.79, p = 0.45). Significant differences also were not detected between ISO and SEP in any of the individual behaviors (agonistic, body-sniffing, rear-sniffing, face-to-face; paired t-tests, p >> 0.0125; **Fig 2C**). Importantly, while there were no overall differences between these two conditions over the full 20 min period, differences were observed in interaction levels during the start of the session. The differences were most evident between 40 to 70 s, as illustrated in **Fig 2D** and identified quantitatively using a series of paired t-tests, revealing a window of 2 s time bins between 42 and 66 s with a 95% likelihood that the two curves differed. At 60 s, for example, ISO degus had interacted on average 20% of the time, while SEP degus had interacted an average of only about 5% (paired t-test, t_7_ = 2.47, p = 0.043, Cowen’s d = 1.16). There was a corresponding statistical trend for steeper initial slopes in cumulative interaction behavior in the ISO condition compared with more evenly paced interactive behavior in the SEP condition (paired t-test on the latency to half-amplitude, t_7 =_ 2.10, p = 0.074). In summary, although interactive behavior between SEP and ISO conditions did not show a net difference over the 20 min period, interactions tended to take place earlier following isolation (ISO), while being more evenly distributed over the session following only separation (SEP).

To help confirm that overall patterns were not due to the order that conditions were presented, direct comparisons were made between interaction levels in Experiments 1 and 2. Interaction levels in SEP (first exposure) were significantly higher than those of 1 min, Experiment 1 (including only dyads whose first exposure was 1 min isolation: two-sample t-test, t_14_ = 2.35, p = 0.034). Likewise, CTL showed significantly lower interaction levels than those following 24 hr isolation in Experiment 1 (equivalent to ISO; only dyads with 24 hr as the second or final exposure included; t_14_ = 2.28, p = 0.039).

By comparing the temporal relationships between physical interactions in ISO and SEP we found a significant decrease in the proportion of rear-sniffing cases preceded by allogrooming/body-sniffing (paired t-test, t_6_ = 3.71, p = 0.010). Importantly, patterns in ISO replicated those observed in the 24 hr condition of Experiment 1 (**Fig 2E**). In other words, while there were no apparent systematic differences in the total levels of face-to-body contact between conditions, degus who had been previously isolated were more prone to engage in allogrooming (“nuzzling”) immediately prior to rear-sniffing. This may reflect a tendency toward affiliation in animals that had been previously deprived of social stimulation. All other temporal relationships were highly consistent between ISO and SEP, and consistent with those observed in Experiment 1 (e.g., face-to-face contact prior to allogrooming/body sniffing, **Fig 2F**).

As was observed in Experiment 1, vocalization rates were highly variable between sessions and dyads. There were no significant differences across the three conditions (repeated-measures ANOVA, F_2,14_ = 2.27, p = 0.14); however, it seems likely that lower vocalizing in CTL would be observed with more statistical power (paired t-test between ISO and CTL, t_7_ = 2.30, p = 0.055, Cohen’s D = 0.78). Given the low power and high variance, it is also unsurprising that we observed no interaction between condition and vocalization type (2-way ANOVA, effect of condition: F_2,300_ = 3.94, p = 0.020, effect of type: F_14,300_ = 19.07, p = 7.03 x 10^-34^, condition x type: F_28,300_ = 1.25, p = 0.19; **S5**). More generally, however, there appeared to be a higher ratio of non-chitters to chitters in SEP compared with ISO (less than half the number of non-chitters to chitters in ISO, and around twice the number of non-chitters to chitters in SEP; a non-parametric, Wilcoxon rank-sum test was used because ratios did not distribute normally, p = 0.0080). This was notable in that a similar but stronger pattern was observed in strangers (Experiment 4, below).

Our repeated-k-means clustering method revealed only one cluster with different proportional expression in SEP versus ISO conditions (**Fig 2G & H**). The syllable type was most strongly revealed when all syllables were subdivided into 8 clusters (d.f. = 7, median p = 0.038, Cohen’s D = 1.01, across 20 repetitions). Unexpectedly, while the cluster was highly overlapping with the one identified in Experiment 1, and typically labeled as “chaffs”, these syllables were expressed more in SEP than ISO (**Fig 2I**). The syllables of the cluster identified in Experiment 1, contained within this larger grouping, were themselves also more common in SEP than ISO (d.f. = 7, p = 0.037). It is worth noting that in both cases—24 hr isolation in Expeirment 1 and ISO in Experiment 2—this was the first time dyads had been reunited following 24 hrs apart. Comparing ISO and CTL, several non-overlapping clusters reached criteria, but the effect magnitudes were very low relative to the other clusters identified and did not lend themselves to straightforward interpretation. Two of these clusters showed proportionally higher activity in ISO but were distributed across several manually-labeled categories (e.g., chaffs, groans, chirps, tweets, whines; d.f. = 7, median p = 0.023 and p = 0.027), another, of manually labeled tweets and chaffs, was found to be proportionally more common in CTL (median p = 0.027).

In summary, Experiment II confirmed our prediction that adult female degus were highly motivated to interact with one-another even after being only separated, and not isolated, for 24 hours, and also confirmed our prediction that the types of interactions among separated individuals differed from those between isolated animals. SEP reunions included more non-chitter to chitter vocalizations than ISO, and ISO elicited more allogrooming/body sniffing before rear-sniffing (consistent with data from Experiment 1). Exploratory analyses identified a chaff-type vocalization that may have been relatively increased in the separation compared with isolation condition. Counterintuitively, this was effectively the same vocalization as was found to increase with 24 hr isolation in Experiment 1, the one common factor between the two being that both Exp II ISO and Exp I 24 hr conditions were the first time dyads had been reunited after extended (24 hrs) apart.

### Experiment III: Acute Stressor vs. Reward

One factor that might have played a role in the differences between isolation and separation is the stress associated with isolation [17–22]. To better understand the ways in which stress modulates female social behavior, we examined behavior following a short period of exposure to an acute, unpredictable stressor (a series of 10, uncued footshocks spaced over 2.5 minutes within a 45 min isolation period; SHK), as compared against the effects of 45 minute isolation in a neutral environment (NTL), and exposure to an unpredictable reward (un-cued delivery of sunflower seeds over a 2.5 minute period within a 45 min isolation period; RWD; **Fig 3A**).

**Fig 3.**
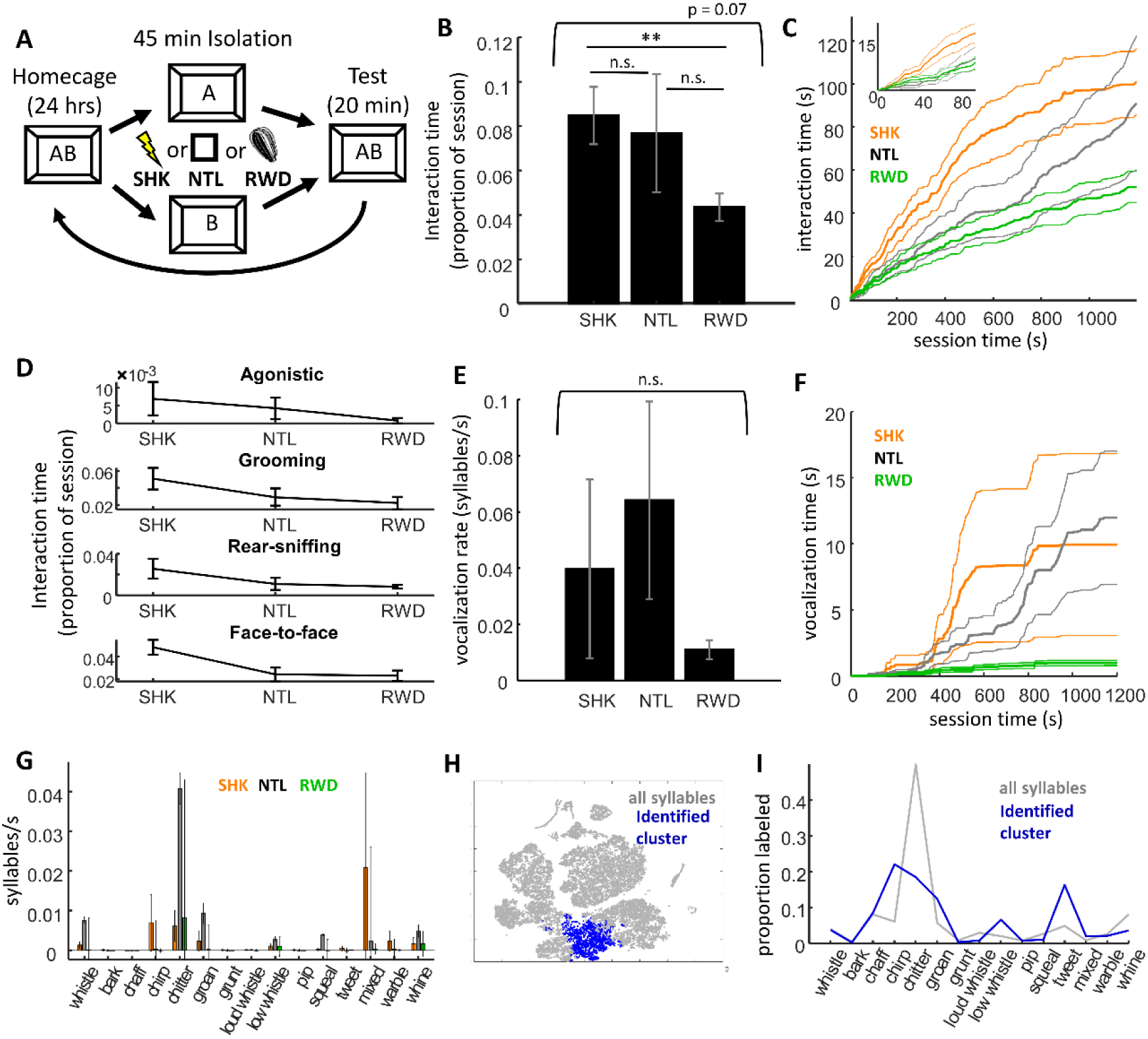
Effects of exposure to an acute stressor on reunion behavior. **A)** Experimental design. Degus (n = 9 dyads) were isolated for a 45 min period during which they were exposed to unexpected footshock (SHK), unexpected reward (sunflower seeds; RWD), or neither (NTL). **B)** Proportion of time dyads spent interacting in each condition. **C)** Cumulative distribution of time spent interacting across the session (thicker lines are means, thin lines standard error). Footshock stressor increased social interaction earlier during the testing, though this behavior leveled off in the latter half. **D)** Four line graphs depict the proportion of time degus spend interacting 10 min into the testing session across each of the three conditions. There was a general pattern for higher interaction across behavior types in SHK at the 10 min mark of the reunion. **E)** Vocalization rates (syllables per second) across conditions. Vocalizations rates were highly variable between dyads. **F)** Cumulative time spent vocalizing (y-axis) is plotted against session time (x-axis). Although there was high variance between dyads, the general pattern was for vocalizations to begin after several minutes of physical interactions, and to subsequently approximately follow the temporal patterns of physical interactions. **G)** Vocalization rates are plotted across each of 15 syllable categories. High variance made interpretation difficult, but generally rates of chitters seemed particularly high in NTL relative to SHK and RWD. **H)** Two dimensional t-SNE projection of syllable features, including those for all syllables (gray points) and those of the cluster of syllables found to be expressed more frequently in SHK. **I)** Proportion of syllables manually labeled across different category types, including all syllables (gray trace) and those syllables within the identified cluster (blue trace). The data suggest that chirp- and tweet-type vocalizations may be more common in degus recently exposed to a footshock stressor.

Physical interaction was affected by a preceding stressor primarily with respect to how the actions were distributed over time. When taking into account the entire 20 minutes, differences between SHK, NTL, and RWD conditions were hinted at by a statistical trend (repeated measures ANOVA, F_2,16_ = 3.93, p = 0.074; **Fig 3B**), although these numbers were largely diluted by the high variance of the neutral 45 min isolation condition, as a clear difference was observed in interaction levels between SHK and RWD (t_8_ = 3.73, p = 0.0058). Looking only at the first half of the session, however, it was clear that the acute stressor did increase initial interactive behavior (**Fig 3C**). Given an alpha of 0.05, there were extended stretches of time in which cumulative interaction time differed between conditions, with differences being maximal between 4 to 6 min (SHK versus NTL, 28 to 100 s and 146 to 958 s, maximum difference observed at 353, t_8_ = 5.30, p = 7.2 x 10^- 4^; SHK versus RWD, from 42 s to the end of the session, maximum at 255, t_8_ = 4.10, p = 0.0013). Paralleling differences between ISO and SEP in Experiment 2, this was accompanied by a statistical trend for steeper initial slopes in the cumulative interaction time (paired t-test on the latency to half-amplitude, t_8_ = 2.15, p = 0.064). There was no evidence that differences between conditions differed between interaction types (2-way ANOVA, effect of condition: F_2,96_ = 3.8, p = 0.026, effect of type: F_3,96_ = 10.13, p = 7.4 x 10^-6^, condition x type: F_6,96_ = 0.48, p = 0.82); however, consistent with the notion that the stressor may have differentially increased affiliative compared with agonistic or investigative behavior, during the first half session SHK sessions included much higher levels of allogrooming/body sniffing and face-to-face contact compared with both NTL (paired t-test, grooming: t_8_ = 4.12, p = 0.0033, face to face: t_8_ = 3.29, p = 0.011) and RWD (grooming: t_8_ = 3.30, p = 0.011, face-to-face: t_8_ = 3.41, p = 0.0092; **Fig 3D**). No differences in the temporal relationships between physical interactions were found between any condition (alpha = 0.05), in all three cases levels of allogrooming-before-rear-sniffing appeared to be intermediate between those observed in ISO/24 hr and SEP conditions.

Similar to Experiments 1 and 2, vocalizations were highly variable across sessions, and no differences were observed between SHK, NTL, and RWD (repeated measures ANOVA on vocalization time: F_2,16_ = 1.24, p = 0.32; **Fig 3E & 3F**). Nor was there a clear differential effect of condition on different types of manually-scored vocalizations (two-way ANOVA, effect of condition: F_2,315_ = 1.85, p = 0.16, effect of type: F_14,315_ = 2.06, p = 0.014, condition x type: F_28,315_ = 1.1, p = 0.34; **Fig 3G**). Repeated-k-means revealed one cluster of syllables differentially increased in SHK relative to NTL (median p across iterations = 0.015, **Fig 3H**) and an almost completely overlapping cluster that increased in SHK relative to RWD (median p = 0.033), while no clusters met criterion when comparing NTL and RWD (alpha = 0.05). These syllables were relatively short in duration (89±14 ms) and were often manually labeled as tweets or chirps (**Fig 3I**).

Summarizing Experiment 3, data confirmed the prediction that footshock stress would increase interaction time, though unexpectedly this was only true for the first half of the session. Likewise, the only common pattern between 24 hr isolation of Experiments I and II and SHK of Experiment 3 was the increased early-session interaction; neither a higher incidence of chitters nor increased allogrooming/body-sniffing before rear-sniffing were observed in SHK. Exploratory analyses identified only a potential increase in tweet/chirp type vocalizations in SHK, and no consistent differences in vocalizations between NTL and RWD

### Experiment IV: Stranger vs. Cagemate

Results from Experiments 1 & 2 were consistent with the possibility that reunion behavior was partly driven by relationship renewal. To further investigate behaviors associated with relationship formation, we compared behavioral patterns in stranger dyads united after 24 hours of isolation (STR) with the the 24 hour cagemate isolation data from Experiments 1 and 2 (CAG1, CAG2; **Fig 4A**).

**Fig 4.**
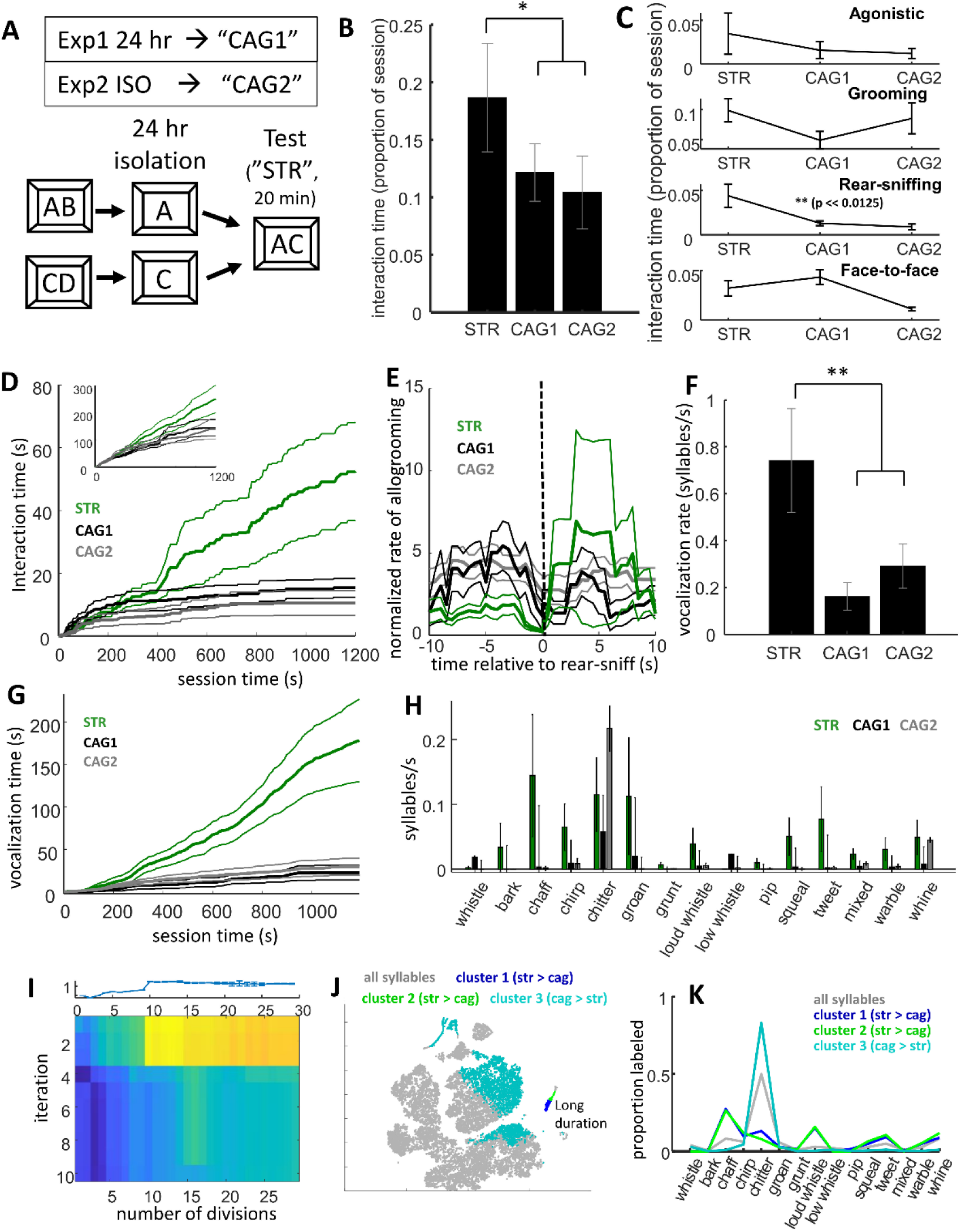
Comparison of social behavior between stranger and cagemate dyads. **A)** Illustration of protocol. Cagemates were isolated for 24 hour and then brought together with an unfamiliar conspecific. **B)** Proportion of time degus spent interacting. There was a statistical trend for stranger pairs to be more interactive than cagemates (STR = stranger, n = 8 dyads; CAG1 = cagemates Experiment 1, n = 9 dyads; CAG2 = cagemates Experiment 2, n = 8 dyads) **C)** Proportion of time dyads of each group engaged in different types of physical interaction. Although there was a pattern for higher levels of interaction between strangers across all types, a strong difference was only observed for rear-sniffing. **D)** Cumulative time dyads in each group spent engaged in rear-sniffing behavior (thicker lines are means, thin lines s.e.m.). Although rear-sniffing was common across all groups during the first several minutes, this behavior decreased among cagemates (CAG1 and CAG2, black and gray traces) but continued at high levels in strangers (STR, green trace). Inset shows cumulative interaction time across all interactive behaviors. **E)** Cross correlogram of levels of allogrooming/body-sniffing (traces) relative to rear-sniffing (see Fig 2E). Similar to SEP in Experiment 2, the STR group showed a lower incidence of allogrooming immediately before rear-sniffing compared with 24 hr isolated cagemates. **F)** Vocalization rates across groups. More vocalizations were observed in STR. G) Cumulative distributions of vocalization time across groups. After a quiet first few minutes, vocalizations were common in STR throughout the session. **H)** Vocalization rates across all 15, manually-labeled syllable categories examined in the present study. Higher vocalizing in strangers was observed across many types, although this was not as clear for chitters (the most common type of syllable observed between cagemates). **I)** Heatmap of identified syllable clusters with the strongest differences between STR and CAG groups (see Figs 1G and 2G). In contrast with prior experiments, the STR versus CAG difference showed multiple independent clusters with differential, proportional activity between the groups, as illustrated by the hot colors observed in each of the 3 top rows. **J)** t-SNE projection of vocalization features (see Figs 1H and 2H). Syllables from each of the three significant clusters are plotted (blue, green, and turquoise). The first two clusters are made-up of particularly long-duration syllables that cluster in a small island; these are most likely combinations of multiple overlapping syllables, notably far more common in stranger dyads than cagemates. **K)** Proportion of syllables manually labeled across different category types (as in Figs 1I, 2I, and 3I). Labeling of the long-duration clusters (blue and green) was variable, most likely because they comprised multiple overlapping syllables between the two individuals. The turquoise trace represented a group of syllables that was proportionally (relative to all syllables) higher in cagemates. These were typically labeled as chitters.

Stranger dyads interacted more than did cagemates (two-sample t-test, n = 8 stranger dyads, 17 pooled cagemate dyads, t_23_ = 2.29, p = 0.032; **Fig 4B**). The effect was largely driven by higher levels of rear-sniffing (anogenital investigation) between strangers (two-sample t-test, t_23_ = 3.55, p = 0.0017, Bonferroni corrected alpha = 0.05/4 = 0.0125; **Fig 4C**). Over the first several minutes of the session interaction levels between STR were on par with those of CAG1 and CAG2; however, while those of cagemates began to decrease and level off, those between strangers continued to remain high. This was particularly evident for rear-sniffing (**Fig 4D**).

Temporal relationships were also different between STR and CAG groups. Paralleling the effects of separation relative to isolation (Experiment 2), strangers were less likely to engage in putative allogrooming immediately before rear sniffing (two-sample t-test, t_23_ = 2.02, p = 0.0075, **Fig 4E; S6**). This result is consistent with the idea that the allogrooming/body-sniffing taking place before anogenital investigation represents an affiliative gesture between animals that had been previously isolated.

Vocalization rates were very high between strangers, exceeding those between cagemates following 24 hours of isolation (two-sample t-test pooling cagemate groups, t_23_ = 3.08, p = 0.0054, **Fig 4F**). Higher levels of vocalizing between strangers was not uniform across syllable types (2- way ANOVA, effect of condition F_2,330_ = 11.5, p = 1.5 x 10^-5^, effect of type: F_14,330_ = 5.33, p = 3.6 x 10^-9^, interaction between condition and type: F_28,330_ = 1.95, p = 3.5 x 10^-3^). Among manually-scored vocalizations, increases appeared particularly pronounced for chaffs (two-sample t-test, pooling cagemate groups, t_23_ = 2.59, p = 0.016), chirps (t_23_ = 2.61, p = 0.016), grunts (t_23_ = 2.88, p = 0.0085), loud whistles (t_23_ = 2.38, p = 0.026), pips (t_23_ = 2.21, p = 0.037), squeals (t_23_ = 2.88, p = 0.0084), tweets (t_23_ = 2.58, p = 0.017), the mixed category (t_23_ = 3.08, p = 0.0054), and warbles (t_23_ = 2.63,p = 0.015). No evidence of difference was observed for chitters, whistles, and whines (p > 0.25). More generally, STR exhibited higher non-chitter to chitter ratios (paralleling SEP and ISO differences in Experiment 2: Wilcoxon rank sum test, p = 0.035). These results were consistent with results of the repeated-k-means method. Three clusters of syllables were identified as being differentially expressed between STR and CAG, but the first two were part of a small cluster of very-long duration (>1 s) syllables that were likely, in fact, combinations of several syllables, and might have combined vocalizations from both individuals (these were often labeled as chaffs, loud whistles, tweets, and whines). The final cluster was comprised of chitters, found to be differentially expressed (relative to all syllables) in CAG. This is consistent with the idea that chitters represented affiliative interactions more prevalent between familiar individuals.

One question that could be raised is whether the differences between strangers and cagemates could be due to kinship differences, as most cagemates were also siblings, while strangers were not born from the same breeding pairs. Since the experiment included only three, non-related cagemate dyads, this could not be thoroughly tested. It is worth noting that non-sibling cagemates, as with siblings, showed lower averages than strangers; however, the effect size for this difference was lower for non-siblings (non-siblings, n = 3, Cohen’s D = 0.51; siblings n = 17, Cohen’s D = 0.94).

In summary, data from Experiment IV confirmed high levels of interaction between strangers, and confirmed that these interactions took a different form than cagemates—including higher rear-sniffing and very high levels of non-chitter vocalizations. Also similar to SEP, strangers did not exhibit high levels of allogrooming/body-sniffing before rear-sniffing observed in cagemates who had been previously isolated for 24 hrs. Exploratory analyses of vocalizations generally confirmed the high degree of non-chitter vocalizing among strangers. Finally, when strangers were compared with non-sibling cagemates effect sizes were reduced suggesting that some differences may have been due to kinship.

## Discussion

### Overview & limitations

The present study was designed to disentangle factors contributing to social motivation in degus after isolation by evaluating details of reunion behavior across different manipulations. A general summary of observations is illustrated in **Fig 5**. Broken-down by experiment, they include the following: **Experiment 1** found that adult female degus become more motivated to interact following longer periods of social isolation. This is a novel result. Although it conforms to observations that have been made in other non-solitary species, including humans [1] and the traditional laboratory model of young adult male rats [2], work in some species has counterintuitively found reduced interactive behavior during reunion (e.g., lion tamarins: [44], and in some cases, marmosets: [45]). **Experiment 2** found that separating degus of a dyad without isolating them from other conspecifics also elicited high interaction levels. Multiple differences were observed between reunions following isolation and separation: after isolation, interactions were distributed earlier in the sessions, included more frequent allogrooming-to-rear-sniffing transitions, and included a higher proportion of chitter vocalizations. Direct comparison of the effects of isolation versus separation-without-isolation have rarely if ever been made between non-human adult, same-sex peers, although results are consistent with other lines of research, including studies on greetings and research opposite-sex, pair-bonded individuals, discussed below. The observed patterns have implications for understanding behaviors dyads may use to relieve isolation stress compared with those they may use to renew previous relationships. Finally, **Experiments 3 and 4** provided further evidence that “stress” and “relationship establishment” are useful constructs for interpreting the behaviors: exposure to a footshock stressor (Experiment 3), like isolation, elicited more early-session reunion interactions, and interactions between strangers contained low levels of pre-rear-sniff allogrooming and relatively higher levels of non-chitter vocalizations. Differences between footshock and isolation suggest stress was not the only factor mediating motivation after isolation; e.g., the apparent nuzzling before rear-sniffing, first identified in Experiment 1 (24 hr vs 1 min) but replicated in Experiment 2 (24 hr ISO vs 1 min CTL), was highest between cagemates after 24 hr isolation, lowest in both strangers and dyads separated without isolation, and apparently at intermediate levels in 45 min isolated animals independent of the stress manipulation performed. The importance of these patterns for understanding relationship renewal as compared with conspecific-general motivations like stress buffering are discussed below.

**Fig 5.**
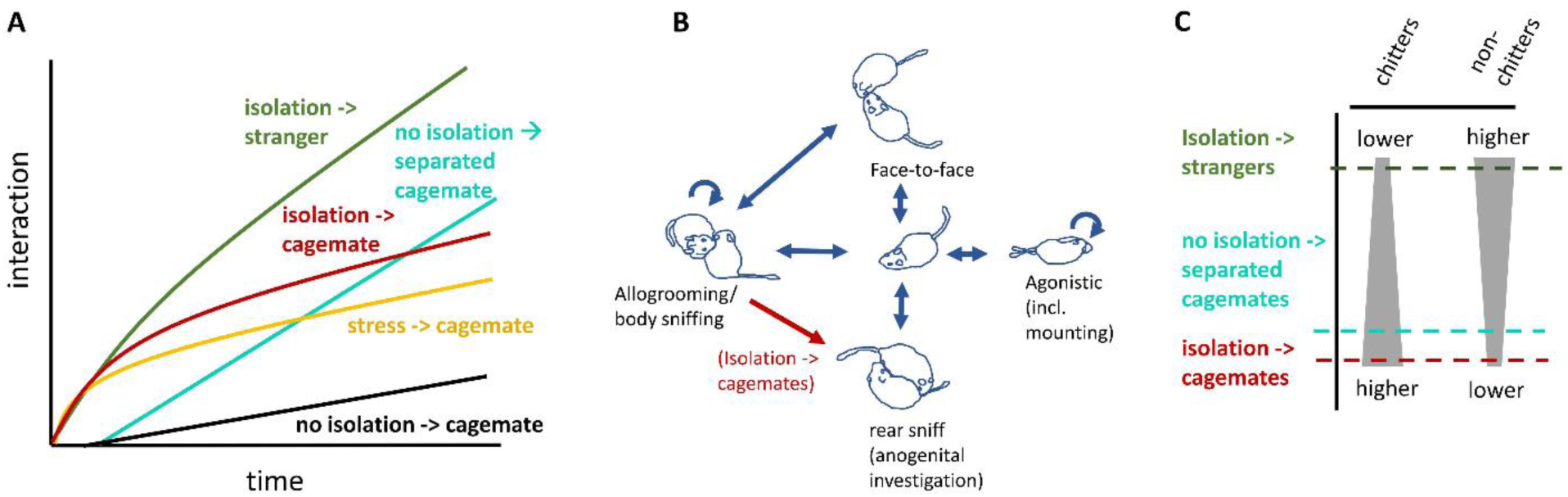
Overview of observed differences between conditions. **A)** Amount of interaction over time (stylized illustration based on data). Both social isolation and footshock stress resulted in higher early-session interaction, while reunions with strangers and previously separated cagemates led to increased interaction over a more prolonged time period. **B)** Illustration of significant transitions between interaction types. An increase in allogrooming before rear-sniffing was primarily observed only in cagemates after isolation. **C)** General differences in vocalizing between groups. Strangers showed highest levels of vocalizations but these were primarily across non-chitters, while the chitter syllable rate in cagemates was very high. Non-isolated, previously separated cagemates showed slightly more resemblance to strangers.

It should be noted that one weakness of the present study is a relative absence of counterbalancing in the order that reunion conditions were presented. In the case of Experiment 3, this was a deliberate strategy to reduce interference of reward and footshock exposure on subsequent reunions, and no progressive changes in social behavior were found across the three conditions. Some of the more salient results of Experiments 1 and 2 were validated by comparing reunion data between the two Experiments (*see Results, Experiment II*). This confirmed higher interaction levels in separation relative to 1 min isolation controls, and lower levels of interaction in 1 min relative to 24 hr isolation. The relatively consistent order of experiments, and conditions within experiments, still raises questions of whether some of the findings could be attributed to condition order. Notably, higher levels of “chaff”-type syllables were observed following the first time degus were reunited after 24 hours in both Experiment 1 isolation and Experiment 2 separation conditions (discussed in more detail below, under *Vocalizations during peer affiliation*), suggesting previous experience may in fact impact dyad behavior during reunion tests. Further research will be required to more thoroughly assess how social behavior is influenced by past experience.

Several factors associated with any study of natural behavior can impact results, including individual differences between animals, and biases or blind spots introduced by the ethograms used. Individuals likely differ in their social tendencies (i.e., “personalities”), and these differences may be further compounded for dyad combinations. Due to the relatively low sample sizes of the present experiments (8 or 9 dyads each), it is likely that effects that do not generalize across most dyads also failed to reach statistical significance, and therefore were not reported. The lower sample sizes may miss behaviors that vary between dyads, but can also be valuable for highlighting the “average behavioral patterns” common across most dyads. Another important factor that should be considered during interpretation is how the ethogram we used may have influenced our specific results. Selecting an ethogram for vocalizations was straightforward, as we were able to follow the extensive examinations of Long [34], and then complemented these with an automated clustering algorithm (our repeated-k-means approach). For physical interactions, behaviors were lumped into four types: agonistic (which included mounting as well as more aggressive behaviors like boxing, hindleg kicks, pushes, bites, etc), non-aggressive nose-to-body contact (allogrooming/body-sniffing), anogenital investigation (rear-sniffing), and face-to-face (which included both nose-nose and nose-mouth contact). It may be that visual inspection of face-to-body contact was insufficient for distinguishing between an affiliative behavior, allogrooming, known to play a role in consolation [46], and an information-seeking behavior, body sniffing. Ultimately, increasing sample sizes to reveal variable effects of the manipulations, and modifying our ethogram to tease-apart issues like grooming versus sniffing, would not have changed the overall conclusions that degus interact heavily with separated cagemates and these differ from stress-related interactions.

### Value of degus for studies of social relationships

Several qualities about degus make this species uniquely suited for investigating reunion behaviors associated with peer relationships. Degus are highly social, potentially sensitive to isolation (previously studied in pups [24]), and there is converging evidence that females in particular are predisposed to make new relationships. Female degus show increased fitness with larger group sizes [47], but there is a high turnover of group membership over time [28] suggesting potential survival benefits to prosocial interactions with new individuals. Female degus are also known to share burrows with multiple, often genetically unrelated peers [25–27,30,31,33,48] (in some cases even cohabitating with unrelated species [48]). As a point of contrast, prairie voles reside with only familiar pair mates or family members [49] and tend to great strangers aggressively [50]. When the two species were directly compared using a partner preference test, prairie voles have been found to spend almost all of their time near or huddling with a familiar partner [51], while degus spend their time near and huddling with partners and strangers, exhibiting no average preference for either [29]. (Mice, meanwhile, tend not huddle with either partners or strangers, and spend much of their time in a non-social neutral chamber [51].) In the present study we confirm that degus interact heavily with strangers, but also show that similar drives may support cagemate interactions following a period of separation. These interactions were distributed across many minutes, suggesting they went beyond an ephemeral “greeting” and involved a prolonged period of re-establishing associations. In the case of strangers, there was also a prolonged period of anogenital investigation, perhaps ensuring subsequent social recognition. One other domain that distinguished new and renewing relationships from isolation and stress was vocal communication.

### Vocalizations during peer affiliation

An intriguing feature of degus is their large repertoire of audible vocalizations [24], and the present study used both manually-scored and automatically-defined syllable categories to identify how vocalizations differed between conditions. Vocalizations were unexpectedly variable between sessions and dyads, far more than physical interactions. Also unexpectedly, while physical interactions began within 10 to 20 s of the reunion, vocal communication was not detected until later, within the first 1 to 2 min (e.g., **Figs 3F & 4G**). It seems unlikely that these patterns were only due to shifts from quieter, undetectable vocalizations, although the amplitude distributions of low-frequency syllables did suggest that some fell outside of the detection range (**Supplementary Fig 1B**). Instead, it may be that vocal communication is a “secondary” quality to their social behavior, used only when an animal feels it’s necessary to modulate another’s behavior while patterning physical investigations and affiliative gestures. This would be consistent with the much higher levels of vocal communication observed in strangers, a situation in which individuals may feel more need to avoid misunderstandings. Consistent with this idea, when rats are devocalized by surgical removal of the larynx, their play behavior more readily escalates into fights [52]. The role of vocalizations for averting misunderstood intentions is much better studied in primates [53]. Grunting in baboons, for example, tends to be used in contexts that may help defuse tense situations, such as during potential conflict or when one female approaches a lower ranked female with an infant [54–57]. These patterns have been conceptualized in game theory terms, with repeated encounters requiring different kinds of information transmitted compared with single encounters [55]. In the present study, chaff-type vocalizations were found to be proportionally higher in relationship establishment/re-establishment conditions across three of the experiments. Long [34] reported chaffs as “[O]ften used when the subject is some distance away from a conspecific, but typically directed toward them. Used by males (7%) and females (93%) of all ages, usually during physical separation of observable conspecific (87.3%), but also when meeting an unfamiliar degu (4.5%).” They further specify the effect of chaffs as being approach, and the likely function as encouraging approach. The present work, which found chaffs to be more common in strangers than cagemates, suggests there more context about the state of an animal’s relationship or recent experiences may be useful for understanding this call.

### Relationship in non-human dyads

The term “social relationship” is often not used in rodent research, although has been defined and applied toward explaining animal behavior [58, 59]. While the term is found only sparsely in the rodent literature, social relationships are studied in multiple forms, including dominance (e.g., mice: [60] rats: [61], marmots: [62], voles: [63], hamsters: [64, 65], degus: [66]), pair bonds (e.g., voles: [67]), and same-sex peer affiliation (voles: [50]). Here we operationally define social relationships as the set of predictive associations (expectations), affiliative bonds, and intra-dyadic “roles” (e.g., dominance roles) that two individuals have for one-another. Research across a range of species has examined the importance of greetings for social bonds and dyadic expectations, although different species may show different emphases. As the temporal distance between two individuals increases, their uncertainty about one-another probably does as well, so greetings may be a means by which that learning can take place to mitigate uncertainty. In some cases this means a relief of tension from potential agonistic or competitiveness interactions (e.g., male baboons [7], black and white colobus monkeys [68], and spider monkeys [5, 69]) while in others this may mean a reaffirmation of affiliative and cooperative bonds (e.g., “high risk” greetings in spotted hyaenas [8] and male guinae baboons [6]). Greetings are less well studied in rodents, but there are a number of exceptions. In marmots, greetings seem to progress behaviorally in ways that arguably go beyond simple social recognition [70], and squirrel greetings vary considerably depending on dominance relationships [71]. Consistent with the present study, greetings between degus in the wild show strong differences depending on whether individuals are from the same or different group areas [48]. Where degu co-residents tended to show face-to-face and mounting behaviors, degus from different areas sometimes exhibiting fighting behaviors and chases. Although the present study does not explore the consequences of greeting behavior in previously-separated degus, it does demonstrate that degus are highly motivated to interact with previously separated individuals over a time period of tens of minutes, even in the absence of other stressors, including isolation. It seems reasonable to infer that the consequences of these interactions include re-establishment of expectations, bonds, and dyadic roles.

One goal of the present study was to examine how social motivation is expressed in adults of a species that does not exhibit pair bonds or same-sex partner preferences, but the results can be considered next to those from research in opposite-sex, pair-bonded animals. The effects of both separations and isolations between pair-bonded partners have been investigated in a number of pair-bonding species (e.g., titi monkeys [72], and both lion tamarins and marmosets [44]), but rarely have the two types of manipulations been directly compared. One exception is a study in Zebra finches, which show much more social interaction with a heterosexual partner after isolation than following separation only, particularly displaying more billing (touching of beaks) and allopreening (allogrooming) [73]. Similarly, marmosets show higher social proximity and potentially more grooming with a novel reproductive partner if they have been previously isolated compared with if they have been kept in social housing [74]. But many of the patterns observed between pair-bonded animals likely generalize to same-sex peers considered in the present work, including the multiple modalities which are used to maintain or re-establish bonds after time apart [75]. As already noted, vocalizations are a common characteristic of peer interactions in general [53], though it seems likely that duetting and courtship songs are more exclusive to either opposite-sex or pair-bonded relationships. These few studies offer some hint at the differences and similarities that may exist in separation versus isolation effects across species and types of relationships, though much more work will be necessary to gain a more generalized understanding the impact of lifespan, species, and social context.

### Social isolation and stress

Reunions between cagemates after both isolation and footshock resulted in more early-session relative to late-session interactions. The similarities between the two conditions could, in principle, be mediated by glucocorticoids resulting from hypothalamic-pituitary-adrenal (HPA) activation [76]. Glucocorticoid responses associated with isolation have been reviewed thoroughly by Hawley et al. [9] for primates, pigs, rats, hamsters, voles and mice (particularly well studied in murine rodents [16–22,77,78]while those in response to footshock have primarily been studied in mice (e.g., [79]) and rats (e.g., [40]). Responses following footshock may not be long-lasting, but if the data in murines generalize to degus then we can expect that in the present experiment they remained elevated across the 10 to 20 min interval between delivery of the footshocks and the start of the reunion session. The increased social interaction may, in turn, have reduced or “buffered” stress levels, a phenomenon that has been studied in a range of species [10–15], including even cows and horses [80, 81]).

While stress may account for similarities between reunions after isolation and footshock, isolation was also distinct from footshock, including higher ratio of chitter vocalizations and more pre-rear-sniff allogrooming. There are many possibilities for the differences. For example, vocalization levels were highly variable following footshock stress, and it may be that degus may be less inclined to make sound if there is a potential external threat in the environment. Another factor may be differences in the levels, duration, or controllability of stress between the two conditions, all of which can differentially impact physiology and behavior [76]. But perhaps the most intriguing explanation for differences between social and non-social stressors is the possibility that other mechanisms, including other signaling systems, are responsible for supporting social motivation following social compared with non-social stress. One candidate is the nonapeptide oxytocin, which has a long-standing reputation as a modulator of social information processing in the brain [82–85]. Although the peptide is not traditionally thought of as a stress hormone, levels of oxytocin have been found to increase not only during social stress, but also following physical stressors, e.g., using restraint or other protocols [86–89] (though not cold exposure [86]). Oxytocin may, in turn, protect an animal against some of the negative behavioral and autonomic effects of social isolation [90]. Another class of signaling molecules that may contribute to social motivation following isolation are opioids [16,23,91] (although it has been argued these play a stronger role in primates [92]). Both oxytocin and endorphins have been linked to social touch, and the value of both molecules for buffering stress—and perhaps loneliness—may implicate these signaling systems in driving social behavior following isolation [23]. These signaling molecules likely interact with specific neurotransmitter systems to mediate behavioral changes. Dopamine neurons in the dorsal raphe nucleus exhibit synaptic changes following isolation and are necessary for increased social behavior following isolation [93]. Serotonin pathways likely also help mediate the motivational salience of social stimuli, as demonstrated not only in rodents, by the importance of serotonin release in the nucleus accumbens [94], but also in species as evolutionarily distant as octopuses [95]. Careful dissection of the contributions these different pathways and signaling systems make to social behavior will offer new insights into the multidimensionality of social motivation.

### Conclusion

The present results provide new information on behaviors associated with relationship renewal, stress, and isolation in a highly social, gregarious rodent. These findings are a valuable lead into understanding how animals in general, and female degus in particular, establish and re-establish expectations about one-another after a period of time, as well as how they overcome the loneliness of isolation during reunion.

## Supporting Information

**S1 Figure.**
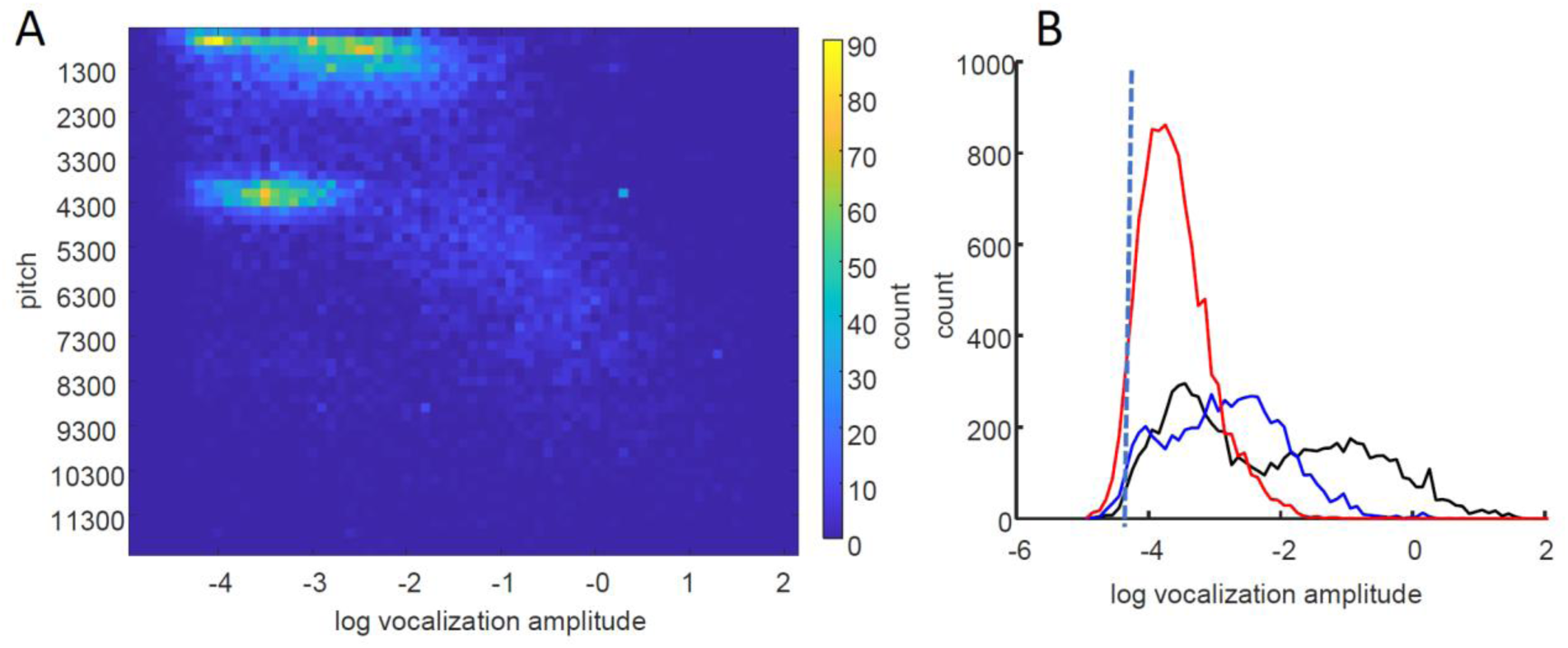
Distribution of maximum amplitude and dominant pitch across all syllables. A) Heatmap of 3D histogram of syllable amplitudes versus dominant pitch. Syllables fell into multiple clusters, the cluster of particularly low amplitudes and pitch were almost exclusively chitters. B) Histogram of clusters that did not have identifiable amplitude peaks above 400 Hz (red trace) and both low pitch (blue trace) and high-pitch (black trace) syllables that did. Black dotted line shows approximates the amplitude threshold for detecting syllables, although this would be more true for the < 400 Hz syllables that fell within the ambient noise. The distribution of these syllables was asymmetric, suggesting that some syllables fell below detection threshold.

**S2 Figure.**
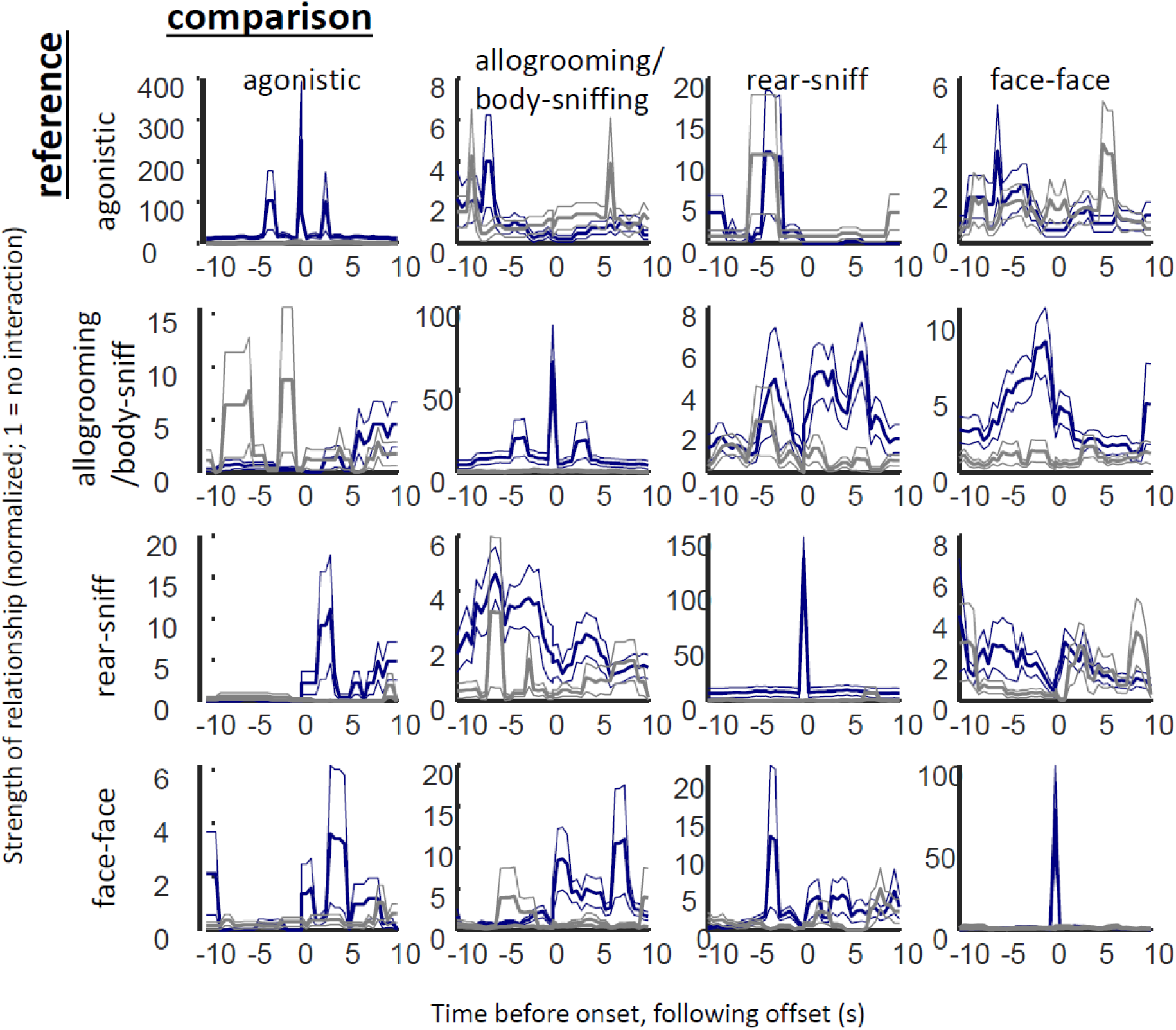
Temporal relationships between physical behavior types. Each plot (blue traces) is a cross correlogram of normalized proportion of time degus spent using a target behavior (columns) relative to a reference behavior (rows). Times before zero are times before the reference behavior starts and times after zero are after the reference behavior ends. Gray traces show the cross correlograms of a surrogate dataset in which times of the comparison behavior are randomly jittered.

**S3 Figure.**
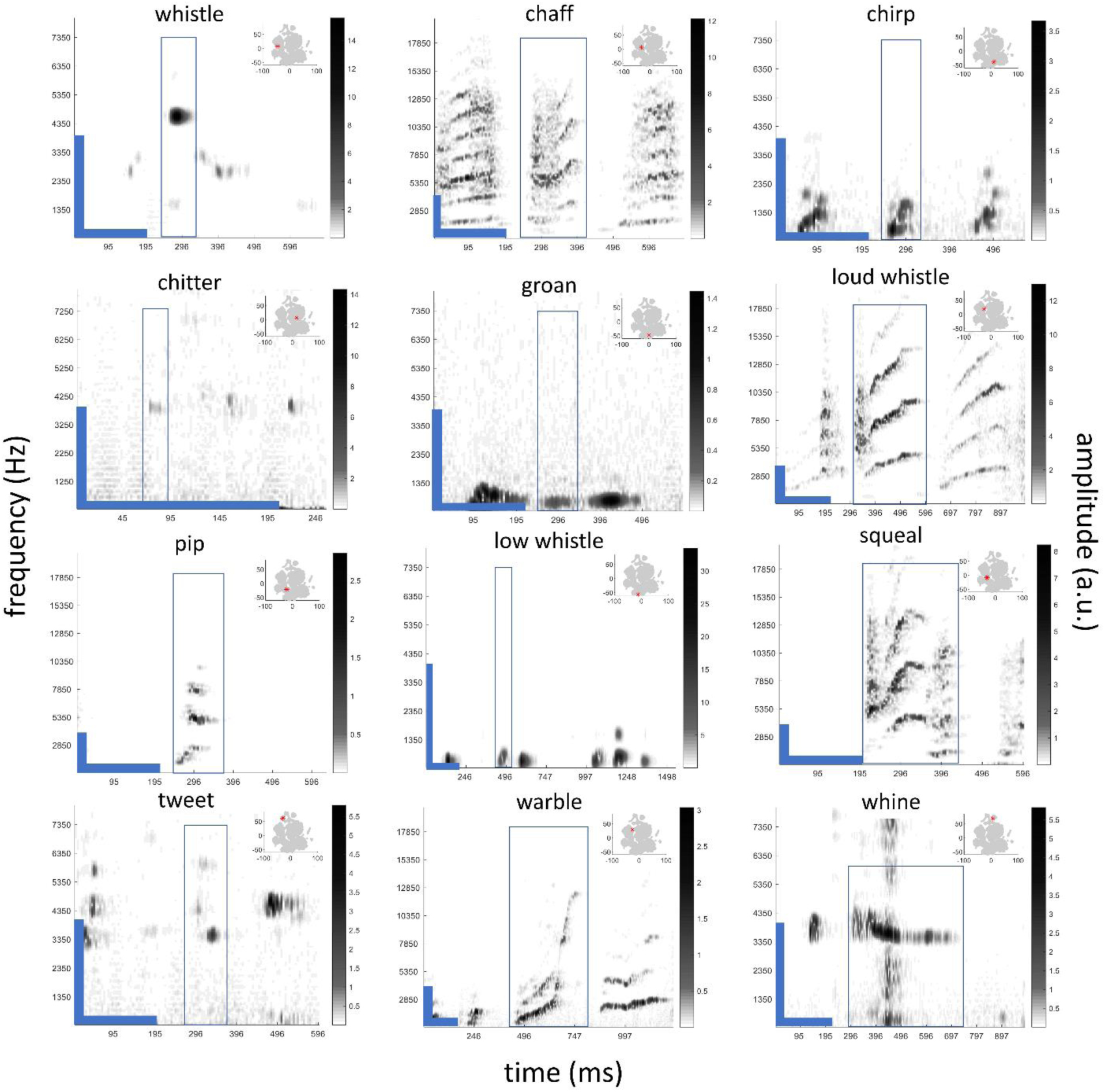
Example spectrograms across twelve scored vocalization categories. Panels show example syllables, with spectral frequencies (y-axis) over time (x-axis) plotted with darker colors reflecting stronger amplitudes. Axes vary across panels: whistle, chirp, chitter, groan, low whistle, tweet, and whines are plotted up to 8000 Hz (y-axis), while chaff, loud whistle, squeal, and warble are plotted up to 20,000 Hz. Variation of y- and y-axes is also illustrated by the blue bars in the lower-left hand corner of each plot (vertical bars indicate frequencies up to 4000, horizontal indicate 200 ms of time), and amplitude strengths with right-side colorbars. Plots begin at 400 Hz to remove low-frequency noise. Box outlines in the center of each plot indicate the specific syllable classified, while time outside of the box is provided for context. Panels in the upper-right show a map syllables in an abstract, two-dimensional projection (using t-SNE, see also **S4**), with all recorded syllables across experiments labeled in gray, and the syllable plotted within the box of the spectrogram identified with a red asterisk. Two syllable categories, barks and grunts are not included because post-hoc examination revealed the syllables were typically misclassified; “mixed” category syllables are also not shown due to the inherent diversity of the category. Black streak in the center of the “whine” example represents background noise.

**S4 Figure.**
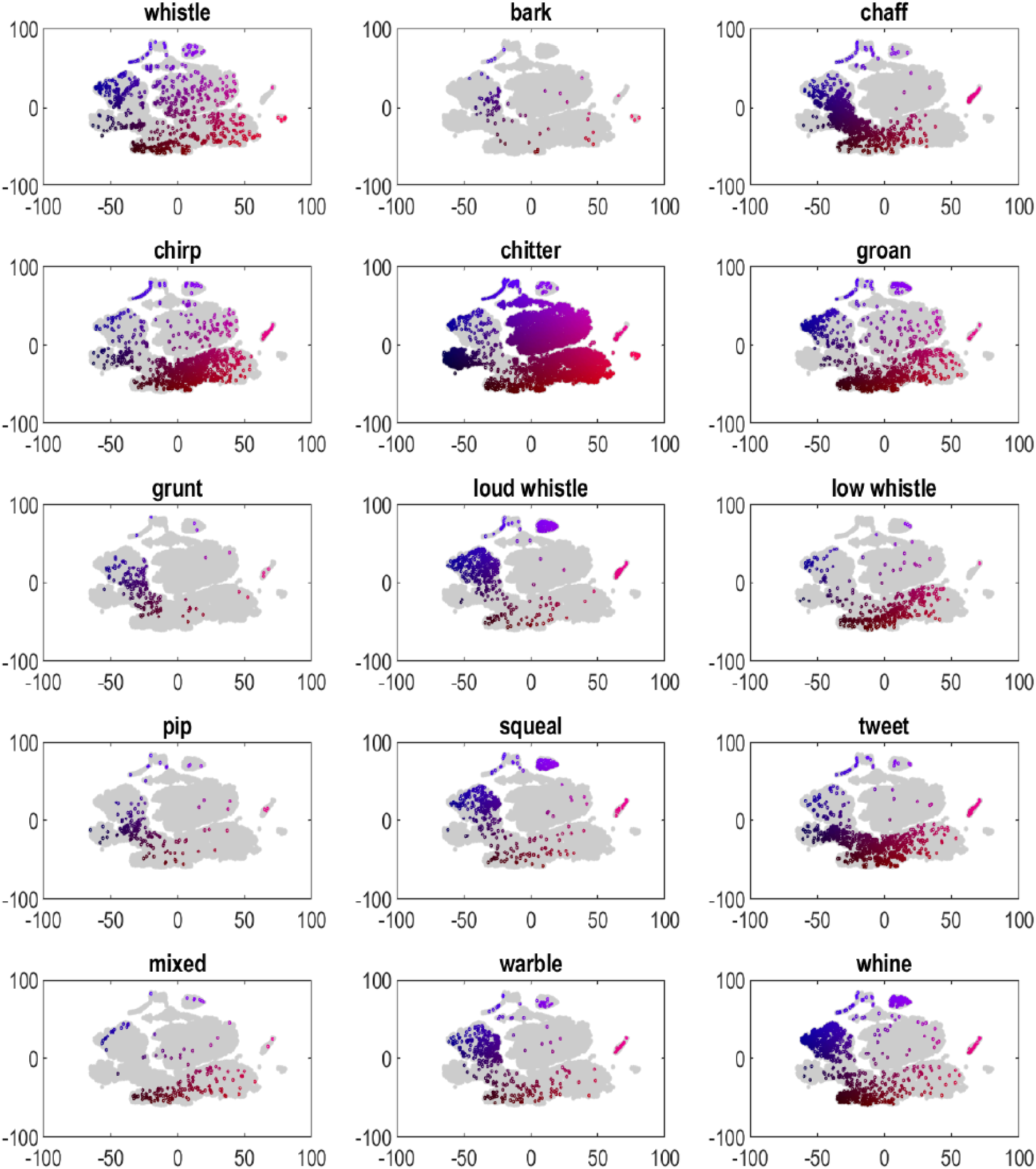
Mapping of labeled syllables to a two-dimensional (t-SNE) projection. Although different vocalization types were distributed differently over multi-dimensional space, there was also a degree of overlap, inspiring the use of automated clustering methods. Axes are in arbitrary units.

**S5 Figure.**
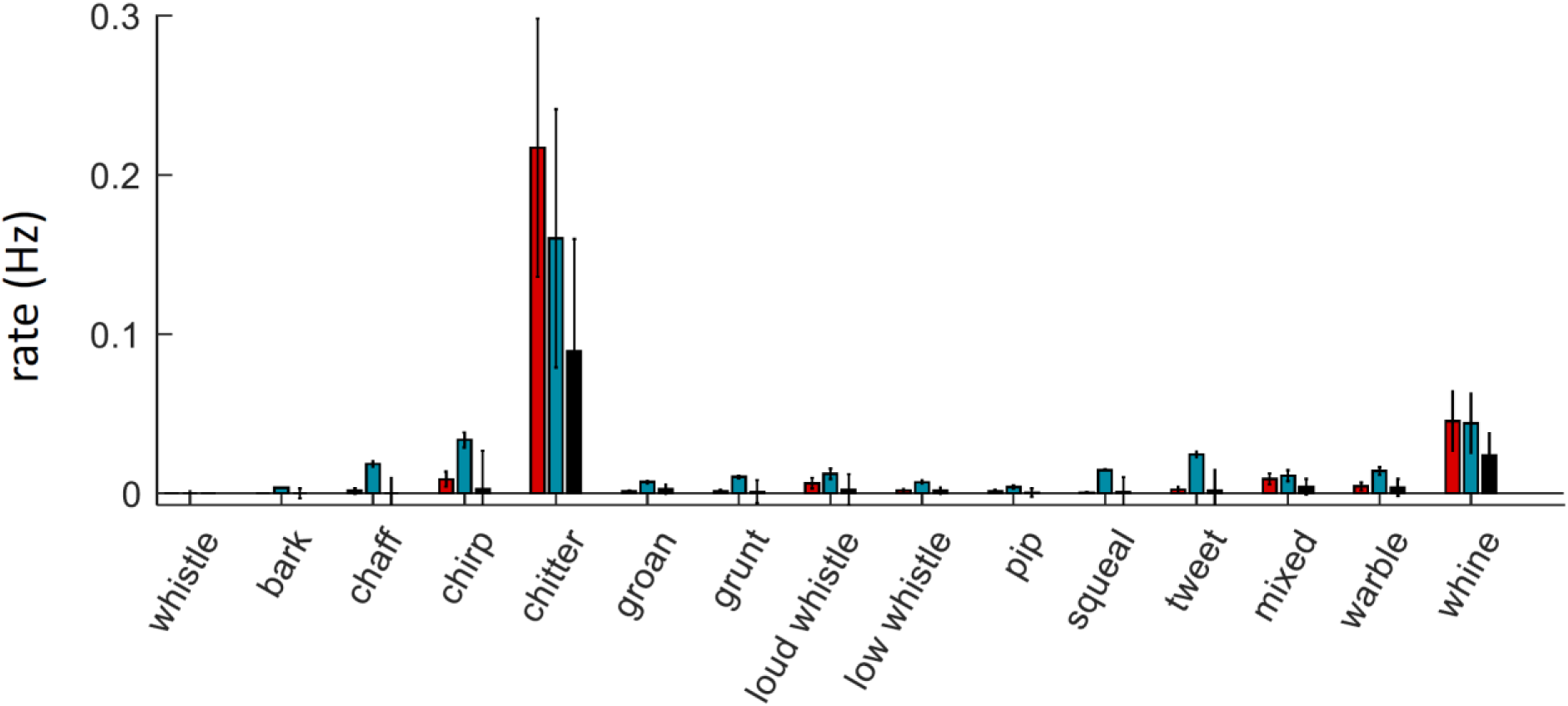
Vocalization rates in Experiment 2. Rates of different vocalization types in the ISO (red), SEP (teal), and CTL conditions (black) conditions. On average, the ratio of non-chitter to chitter vocalizations was higher in SEP.

**S6.**
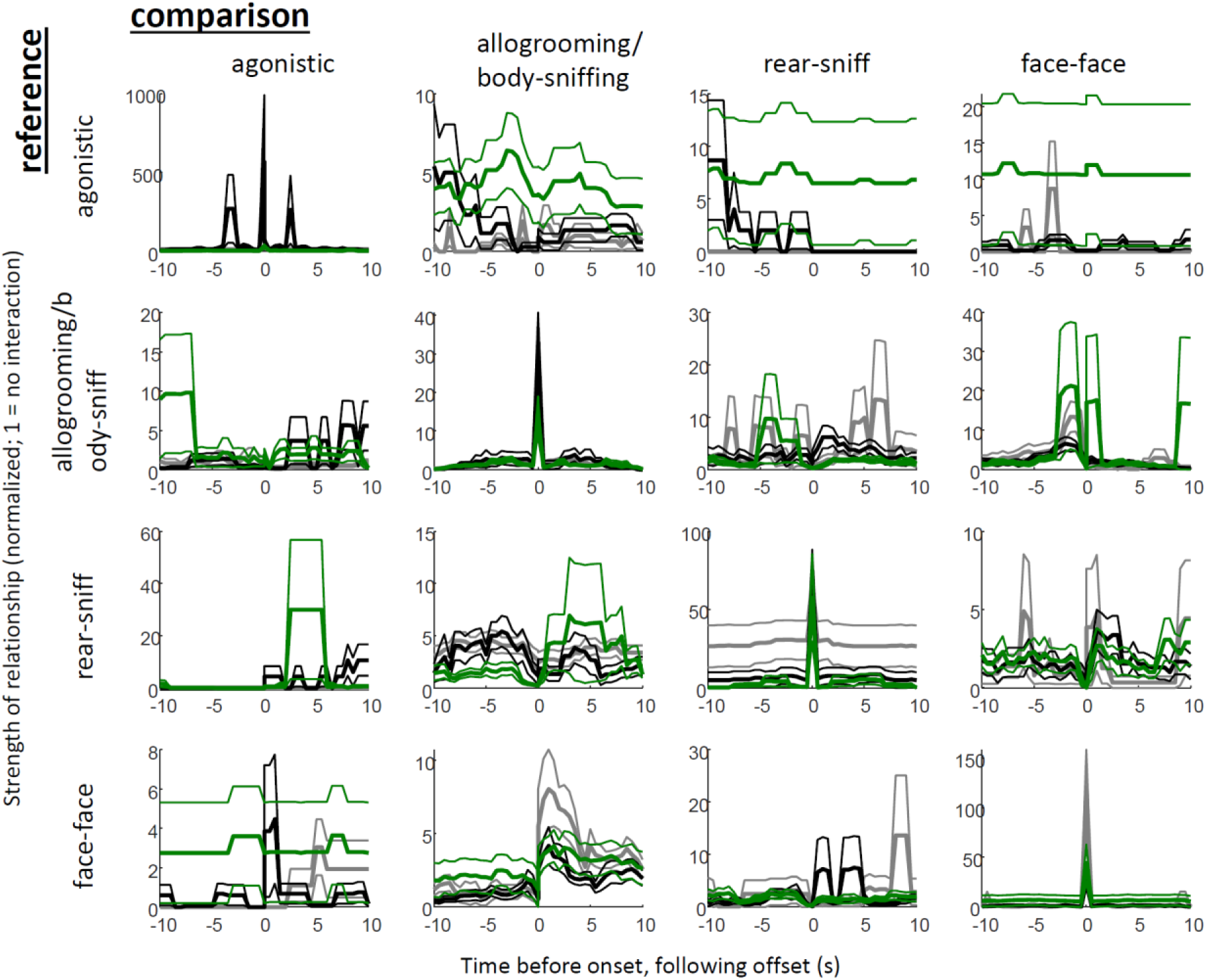
Temporal relationships between interactions in STR and CAG. Green traces are STR, black are CAG1, and gray are CAG2 of Experiment 4, organized as in S2. The only significant difference was observed in the higher levels of allogrooming/body-sniffing that preceded rear sniffing in both CAG1 and CAG2 conditions. Though not statistically significant, the figure also suggests apparently higher grooming/body-sniffing before and after agonistic interactions in strangers; this may have been due to either bites to the body, or overly-vigorous (painful) grooming labeled as agonistic in one or a few stranger dyads.

## Notes

### Competing Interest Statement

The authors have declared no competing interest.

### Summary of Updates

Minor corrections in the table and figures, minor changes in wording and organization of Discussion.

